# G-quadruplex stabilization induces DNA breaks in pericentromeric repetitive DNA sequences in B lymphocytes

**DOI:** 10.1101/2025.05.09.653134

**Authors:** Irina Waisertreiger, Kalkidan Ayele, Mehad Hilal Elshaikh, Jacqueline H. Barlow

**Author notes:** Jacqueline H. Barlow, **Email:**. **Author Contributions:** IW and JHB designed the study, interpreted and discussed all the results, and wrote the manuscript. IW supervised all experiments and performed the bulk of experiments including all human cell work. KA performed and analyzed PDS-induced genome instability alone and in combination with WEE1, and TUNEL experiments. MHE performed and analyzed WEE1 TUNEL, CH12 cell cycle analysis by flow cytometry, and protein blot experiments. KA and MHE assisted with mouse genotyping, and cell culture. IW and JHB led the entire study and JHB obtained funding for the study. All authors contributed to the article and approved the submitted version. **Competing Interest Statement:** The authors declare no competing financial interests.

## Abstract

DNA secondary G-quadruplex (G4) structures can impair and even obstruct the DNA replication. Defects in processing G4 structures are associated with replication stress, a common property of both B cell cancers and hyperproliferative premalignant cells. Genome instability arising from replication stress is a hallmark of cancer and strongly contributes to the chromosome rearrangements in B cell cancers. Here, we define the impact of G4-stabilizing ligands on generating genome instability in primary and malignant B cells. Treatment with the G4-stabilizing compound pyridostatin (PDS) causes breaks and chromosome rearrangements at ribosomal DNA and pericentromeric major satellite regions in both mouse primary B cell culture and CH12 lymphoma cells. PDS also causes extensive pericentromeric DNA damage in immortalized human B cell lines. Remarkably, PDS causes high level of tetraploid metaphase cells correlated with high level of dicentric chromosomes specifically in primary but not in CH12 B cells. Unlike primary B cells, CH12 cells undergo checkpoint activation and strong G2/M arrest in response to PDS treatment thus preventing tetraploid appearance. Altogether, these results highlight the difference between primary and malignant B cells in response to PDS, revealing the therapeutic potential of G4-stabilizing drugs to selectively suppress tumor cell growth and proliferation.

**Significance Statement:** G-quadruplexes are guanine-rich DNA secondary structures, abundantly found in most eukaryotic genomes. Stabilization of G-quadruplexes creates an obstacle for DNA replication, particularly in DNA repair-deficient cells. We show that G-quadruplex stabilization by the small molecule pyridostatin leads to recurrent DNA breaks and aberrant chromosomal fusions in the repetitive DNA sequences found at major satellite pericentromeric repeats and ribosomal DNA arrays in mouse primary B cells. Pyridostatin also causes extensive pericentromeric DNA damage and pathogenic rearrangements in mouse and human B cell lymphoma cell lines. Thus we have found a role for G-quadruplexes in inciting damage driving the formation of recurrent rearrangements observed in lymphoid cancers.

## Introduction

G-quadruplex structures are four-stranded secondary DNA structures that can form in guanine-rich DNA sequences. Guanines stack into a stable planar quartet stabilized by Hoogstein hydrogen bonding, potentially leaving the complementary sequence single stranded (1). Using a variety of prediction tools, over 350,000 sequences with G4-forming potential have been identified in the human genome (2, 3). G4 formation has been more directly mapped genome-wide utilizing G4-specific antibodies or a G4 binding protein to directly identify structure-specific motifs, indicating that thousands of loci exhibit G4-forming capability *in vivo* (4–6). G4s are enriched at regulatory regions of genes and within origins of replication (7–10). These results reinforce prior work indicating roles for G4 structures in regulating transcription as well as replication licensing and initiation (11–14).

To functionally dissect the role of G4 structures *in vivo,* multiple G4-binding ligands have been developed with a wide range of selectivity and *in vivo* effects (15). Pyridostatin (PDS) is a potent and selective G4 stabilizer, thought to interact in a planar manner at the quadruplex/duplex DNA junction stabilized by ν-ν stacking of the aromatic rings (16–18). A major effect of exposure to PDS is cytotoxicity and genome instability (19, 20). PDS-induced cell death is enhanced in cell lines mutated for DNA repair factors associated with homologous directed repair (HDR), suggesting genome instability drives cytotoxicity. Human cells harboring mutations in *BRCA1* or *BRCA2* are particularly sensitive to PDS as well as CX-3543 and CX-5461, two independently identified G4-stabilizing compounds, indicating chemotherapeutic potential for G4 ligands (21–23).

G4 ligand-mediated instability is strongly linked to DNA replication, indicating that G4 structures represent a barrier for replication fork progression (19). Over a dozen helicases are reported to have G4 unwinding capability, and loss of function can induce defects in replication fork progression and instability (24–28). The link between G4s and replication-induced damage is further supported by reports indicating that G4 replication requires an accessory helicase (29). However G4s may not act alone to perturb replication, as G4 formation and damage also correlate with R loop formation (30, 31). PDS was shown to promote telomeric chromosomal instability in some cancer cell lines (22), however studies evaluating the effect of PDS on overall chromosomal stability are limited.

In the current study, we used mouse and human primary B cells and B cell lymphoma cell lines to investigate the genomic location and type of chromosome damage induced by the G4 ligand PDS. We observed PDS-induced chromosomal instability in all examined cell types except human primary B cells. More than half of aberrations involved pericentromeric chromosomal DNA. Mouse cells exhibited PDS-induced DNA breaks at the ribosomal DNA (rDNA) repeats and centromeric major satellite (MaSat) region. Intriguingly, PDS also induced tetraploidization visible in mouse primary B cells, CH12 lymphoma B cells, and human EBV-immortalized GM12878 cells. In mouse cells, tetraploid metaphase cells correlated with a lack of Wee1-dependent G2/M checkpoint arrest indicating mitotic slippage, and artificially enforcing G2 arrest with CDK1 inhibition strongly reduced tetraploid metaphase cells. In contrast, PDS induced a strong G2 arrest in CH12 cells which was reversed by Wee1 inhibition. Wee1 inhibition in CH12 cells also induced tetraploid metaphases. Our data point to tetraploidization as an unexpected consequence of PDS treatment in lymphoid cells, likely dependent on absence of G2/M arrest.

## Results

### PDS exposure causes moderate cytotoxicity in mouse primary B cells

PDS is cytotoxic to mouse and human cell lines, therefore we set out to define the impact of PDS exposure on mouse primary B cells. *Brca1*-mutant cells are particularly sensitive to G4 ligands (21, 22, 32), therefore we exposed activated, proliferating primary B cells expressing either full-length or a mutant form of *Brca1* lacking exon 11—hereafter referred to as *Brca1^111/111^*—to multiple PDS doses for 48 hours. *Brca1^111/111^* B cells are generated by conditional deletion of exon 11 specifically in B cells by co-expression of cre recombinase from the *Cd19* promoter (33, 34). PDS treatment reduced viability in both WT and *Brca1^111/111^* cells (Fig. S1A), and increased apoptosis (Fig. S1B) in a dose-dependent manner. Notably, we observed only a modest difference in cytotoxicity between WT and *Brca1^111/111^* cells at 5 µM. Analysis of the proliferation rate by CFSE-staining showed that 5 µM PDS moderately slowed cell division (Fig. S1C) in both WT and *Brca1^111/111^* cells.

### PDS causes specific chromosomal rearrangements in mouse primary B cells

To test if PDS causes persistent chromosomal instability, we treated WT and *Brca1^111/111^* B cells with PDS for 48 hours and examined metaphase chromosome spreads for gross chromosome rearrangements (Fig. 1A,B). Normal mouse chromosomes are all acrocentric: the extremely short p arms encode for no genes therefore the heterochromatic centromeric region, visible by strong DAPI signal, appears on one end and co-localizes with telomeric probes (Fig. 1A, first column). We found that PDS induces chromosomal instability in a dose-dependent manner, and *Brca1^111/111^* cells consistently show more chromosome aberrations than WT cells; the trend is observable at all doses examined, however the increase was only significant at 5 µM PDS, and the high cytotoxicity at 10 µM likely obscured differences in damage levels (Fig. 1C). This difference in chromosome rearrangement frequency between WT and *Brca1^111/111^* B cells may reflect impaired repair in *Brca1^111/111^* cells, as HDR is important for resolution of G4 stabilization induced DNA damage (22, 35). Interestingly, PDS exposure creates specific types of chromosomal rearrangements in primary mouse B cells: we observed frequent chromosome breaks and two-arm fusions, but few chromatid breaks (Fig. 1A,B,D). Most fusions were dicentrics, Robertsonian chromosomes, two-arm fusions, and acentric chromosomes (Fig. 1A, B). These types of fusions involve two non-homologous chromosomes fused at similar regions on both arms. To test if this was specific to primary cells, we next exposed the immortalized mouse B cell lymphoma line CH12 (36) to PDS and observed a mild increase in apoptosis compared to untreated cells (Fig. S1D). PDS also caused chromosomal instability in CH12 cells (Fig. S1E). Notably, CH12 cells exhibited a similar spectrum of rearrangements as primary B cells; though CH12 cells harbored fewer dicentric fusions (Fig. 1E, Fig. S1F).

**Figure 1.**
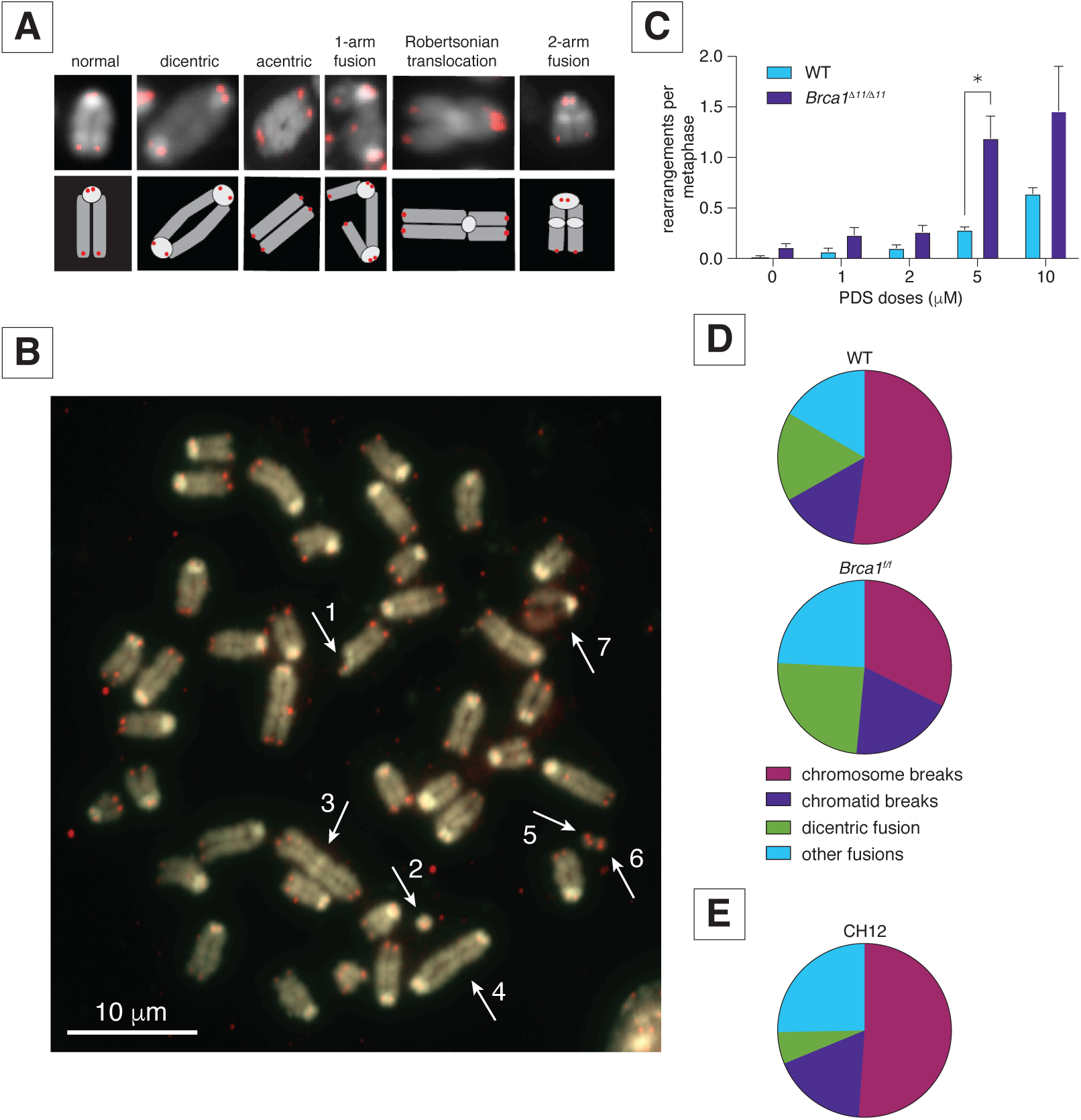
WT, *Brca1^111/111^*, and CH12 cells exhibit specific chromosomal rearrangements in response to PDS. **A)** Representative images (top) and cartoon diagrams (lower) of the specific types of chromosomal fusions produced by PDS exposure. Telomere-specific probe visualized in red, DAPI is in greyscale. **B)** Representative image of metaphase spread with many of the typical chromosomal rearrangements observed in response to PDS. Arrows highlight the rearrangements: #1 – pericentromeric chromatid break; #2 - tiny dicentric chromosome; #3 – acentric chromosome with heterochromatic (probably pericentromeric) block in the middle; #4 – dicentric chromosome; #5 and #6 – fragments from chromosome breaks, probably from the chromosomes formed dicentric #4; #7 – chromosome likely from a different metaphase plate, which shows different chromatin condensation level and significant loss of cohesion between chromatids comparing to other chromosomes. Scale bar – 10 µm. **C)** Number of DNA aberrations per metaphase in response to PDS in WT and *Brca1^111/111^* cells. Error bars show the standard error of mean (SEM) from three independent experiments. Statistics: *p < 0.05 comparing WT and *Brca1^111/111^* cells exposed to PDS. **D)** Fraction of specific rearrangements in response to PDS in WT and *Brca1^111/111^* cells. **E)** Fraction of specific rearrangements in response to PDS in CH12 cells.

### The majority of PDS-induced rearrangements occur at rDNA and MaSat repeats in mouse primary B cells

Unlike prior studies in immortalized mouse and human cancer cell lines, we did not observe PDS-induced telomeric fragility, previously described as either multiple FISH probe signals at a single telomere or loss of telomere signal (22, 23). However we observed that the majority of PDS-induced chromosomal breaks were at or near DAPI-bright centromeric heterochromatin (Fig.1B, arrow #1; Fig. 2A and Fig. 2B, top and middle rows). We also observed a high frequency of very small dicentrics that appeared mostly heterochromatic, suggesting they arise from fusion of pericentromeric chromosomal breaks from two chromosomes (Fig. 1A, arrow #2; Fig. 2A, bottom panel; Fig. 2B, third panel from top). Tandem arrays of rDNA repeats are located near the centromere of chromosomes 12,15, 16, 18 and 19 in mouse and form G4 structures in multiple species, therefore we investigated if rDNA repeats are involved in PDS-induced rearrangements (37–44). Using a BAC probe spanning the entire 45s rDNA region, we found that simple breaks and chromosome fusions at or near rDNA were frequent events in response to PDS (Fig. 2A, C, D). Overall, ∼13.5% of total rearrangements in WT B cells and ∼9.5% of rearrangements in *Brca1^111/111^* cells involve rDNA repeats (Fig. 2G).

**Figure 2.**
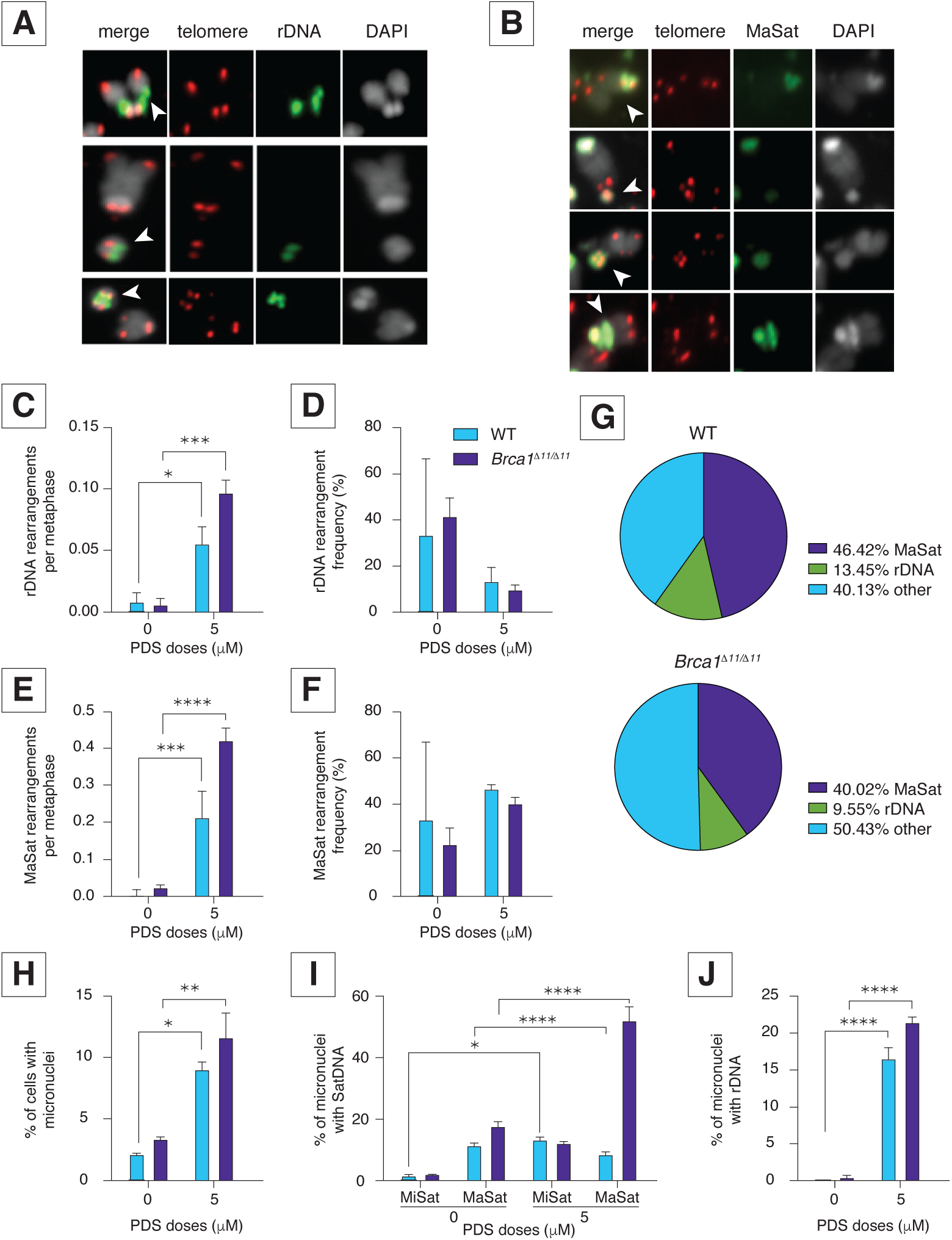
The majority of rearrangements occur at rDNA and MaSat in WT and *Brca1^111/111^* cells in response to PDS. **A)** Representative images of rearrangements in rDNA in response to PDS. Top panel – chromatid break at rDNA. The rDNA probe flanks both break ends. Middle panel - chromosome break at rDNA. Bottom panel – dicentric chromosome with fusion at rDNA. Arrows highlight the rearrangements. Telomere-specific probe visualized in red, rDNA is in green, DAPI is in greyscale. **B)** Representative images of rearrangements in MaSat in response to PDS. Top panel – chromatid break at MaSat. Second from top panel - chromosome break at MaSat. Third from the top panel – dicentric chromosome with fusion at MaSat. Bottom panel – two-arms fusion at MaSat. Arrows highlight the rearrangements. Telomere-specific probe visualized in red, MaSat is in green, DAPI is in greyscale. **C)** Rearrangements at rDNA per metaphase in response to PDS in WT and *Brca1^111/111^* cells. Error bars show SEM from three independent experiments. Statistics: *p < 0.05 and ***p<0.005 comparing PDS doses. **D)** Frequency of rearrangements at rDNA in response to PDS in WT and *Brca1^111/111^* cells. Error bars show the SEM from three independent experiments. **E)** Rearrangements at MaSat in response to PDS in WT and *Brca1^111/111^* cells. Error bars show the SEM from three independent experiments. Statistics: ***p < 0.005 and ****p<0.001 comparing PDS doses. **F)** Frequency of rearrangements at MaSat in response to PDS in WT and *Brca1^111/111^* cells. **G)** Fraction of DNA rearrangements at rDNA and MaSat in response to 5 µM PDS in WT and *Brca1^111/111^*cells. **H)** Percent of cells with micronuclei in response to PDS in WT and *Brca1^111/111^* cells. Error bars show the SEM from three independent experiments. Statistics: **p < 0.01 and *p <0.05 comparing PDS doses. **I)** Percent of micronuclei containing MiSat and MaSat in response to PDS in WT and *Brca1^111/111^* cells. Error bars show the SEM from three independent experiments. Statistics: ****p < 0.001 and *p <0.05 comparing PDS doses. **J)** Percent of micronuclei containing rDNA in response to PDS in WT and *Brca1^111/111^* cells. Error bars show the SEM from three independent experiments. Statistics: ****p < 0.001 comparing PDS doses. For all graphs, WT is turquoise and *Brca1^111/111^* is royal blue.

A significant portion of pericentric aberrations did not co-localize with rDNA, suggesting additional hotspots of fragility. The mouse pericentromeric MaSat (Fig. S2A) also contains a weak G4 forming motif using QGRS Mapper and G4 Hunter (45, 46) (Table S1). Like rDNA, MaSat sequences (Fig. S2A) were involved in both simple breaks and fusion events with ∼46.5% of total rearrangements in WT and ∼40% of total rearrangements in *Brca1^111/111^* cells (Fig. 2B,E-G). Remarkably, we found almost no centromeric minor satellite (MiSat) involvement in PDS-induced rearrangements (Fig. S2A). We observed no chromosome or chromatid breaks flanking the MiSat probe, and most fusions arising from centromere breaks exhibited complete loss of MiSat signal, suggesting that centromeric breaks and consequent fusions involve MaSat, but not MiSat sequences (Fig. S2B, C). We conclude that pericentromeric MaSat but not centromeric MiSat repeats are sensitive to G4 stabilization by PDS.

G4 stabilization induces micronuclei formation as a hallmark of genome instability in cancer (21, 31, 47, 48). We also found an elevated frequency of micronuclei formation in response to PDS in primary B cells (Fig. 2H). Less than 20% of micronuclei contained MiSat signal (Fig. 2I), whereas ∼50% of micronuclei contained MaSat signal (Fig. 2I), and more than 15% of micronuclei contained rDNA signal (Fig. 2J). Taken together, the data indicates that pericentromeric MaSat and rDNA repeats are major locations of PDS-induced chromosomal instability.

### PDS induces high levels of tetraploid metaphase cells in primary but not immortalized B cells

We observed high levels of tetraploidy in PDS-treated cells. Over 10% of WT and ∼20% of *Brca1^111/111^* metaphase spreads were tetraploid (∼78-82 chromosomes) after 5 µM PDS treatment for 48 hours (Fig. 3A, S3A). To determine if we could visualize tetraploid cells prior to mitosis, we performed FISH on interphase nuclei with a locus-specific BCL2 BAC probe. This method allows differentiation between normal diploid cells in G2 from tetraploid cells: Diploid G1 cells exhibit two single (unreplicated) dots while G2 cells exhibit two twin (replicated) BAC signals (Fig. S3B, top two panels). In contrast, tetraploid nuclei have four or more separate FISH signals as either single (unreplicated) or twin (replicated) dots depending on replication status (Fig. S3B, lower two panels). We found similar levels of tetraploid interphase nuclei and tetraploid metaphase cells in WT and *Brca1^111/111^* cells exposed to PDS (Fig. 3B, Fig S3C), suggesting that tetraploids do not experience G2/M cell cycle arrest. Strikingly, we almost did not find tetraploid metaphases in PDS-treated CH12 cells (Fig. 3A), but observed high levels of tetraploid interphase nuclei (Fig. 3B, Fig. S3C). Taken together, these data suggest that PDS can induce tetraploidy in both primary and cancerous CH12 mouse B cells; however tetraploid CH12 cells experience G2/M arrest while tetraploid primary B cells proceed into mitosis.

**Figure 3.**
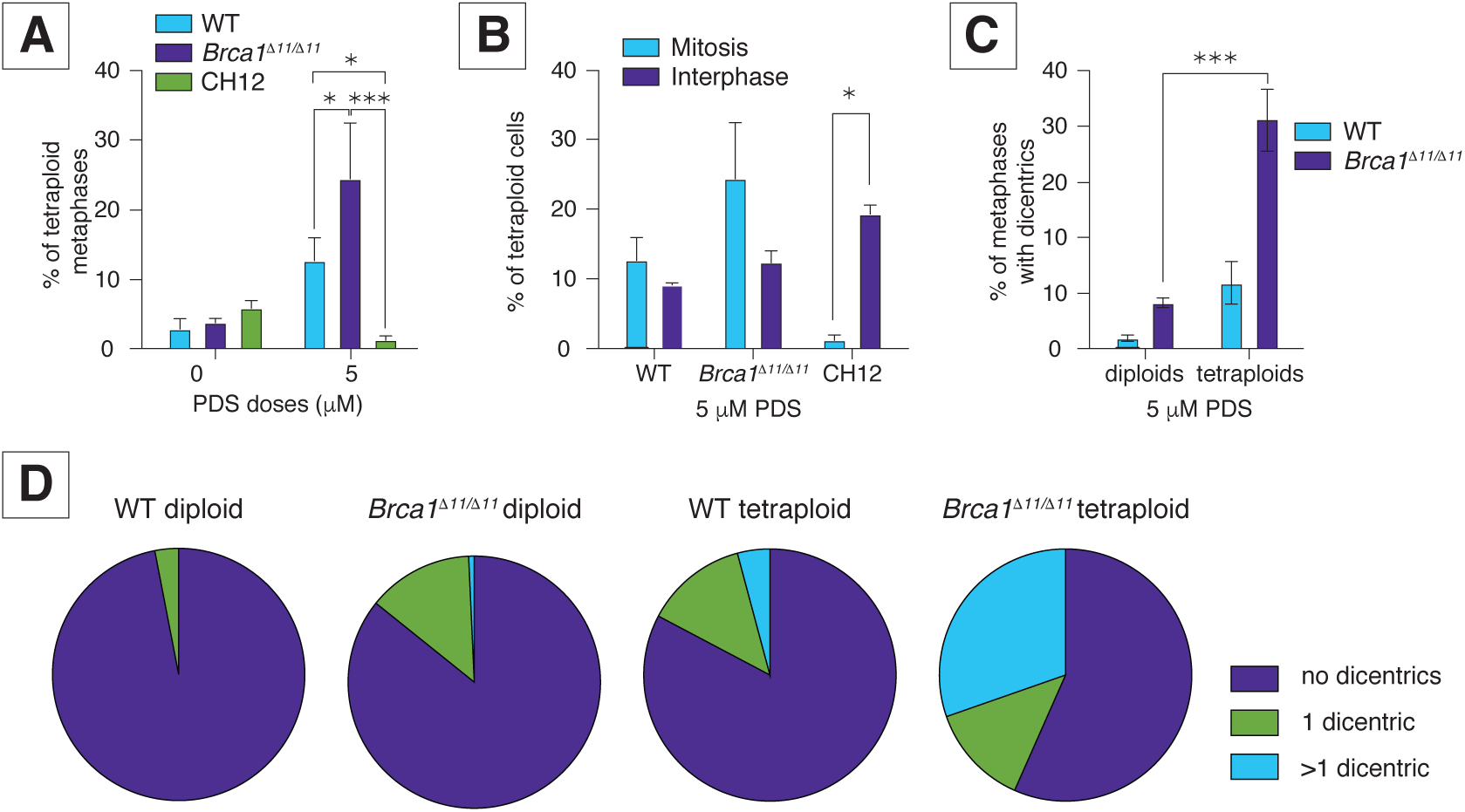
Primary B cells, but not CH12 cells, show high levels of tetraploid metaphases in response to PDS. **A)** Frequency of tetraploid metaphases in response to PDS in WT, *Brca1^111/111^*, and CH12 cells. Error bars show SEM from three independent experiments. Statistics: ***p < 0.005 and *p < 0.05 comparing WT, *Brca1^111/111^*, and CH12 cells exposed to PDS. **B)** Percent of tetraploid cells in mitosis (metaphases) and tetraploid interphase nuclei in WT, *Brca1^111/111^*, and CH12 cells exposed to 5 µM PDS. Error bars show SEM from three independent experiments. Statistics: *p < 0.05 comparing of tetraploid metaphases and tetraploid interphase nuclei. **C)** Percent of diploid and tetraploid metaphases containing dicentric chromosomes in response to 5 µM PDS in WT and *Brca1^111/111^* cells. Error bars show the SEM from three independent experiments. Statistics: ***p < 0.005 comparing diploid and tetraploid cells exposed to 5 µM PDS. **D)** Fraction of metaphases containing no dicentric chromosomes, one dicentric chromosome, or more than one dicentric chromosome cells in response to 5 µM PDS in WT and *Brca1^111/111^* cells.

Tetraploidy can be caused by cell fusion (49), endoreplication (50, 51) or mitotic slippage (52–54). Notably, the tetraploidy observed in mouse primary B cells strongly correlated with dicentrics appearance (Fig. 3C, Table S2). Dicentric and multi-centromeric chromosomes form multi-polar attachments in mitosis, potentially triggering an anaphase spindle checkpoint (55, 56). Moreover, tetraploid metaphases usually carried more than one dicentric chromosome (Fig. 3D, Table S2). It is believed that prolonged mitosis can trigger mitotic slippage (54, 57). We propose that PDS exposure induces tetraploidy in mouse primary B cells by creating multiple dicentrics or other multi-centromere chromosome rearrangements that trigger spindle checkpoint activation and anaphase delay leading to mitotic slippage.

### PDS-induced tetraploid metaphases in primary B cells are associated with absence of G2/M arrest

To test our hypothesis that tetraploid metaphases in PDS-treated CH12 cells were absent due to cell cycle arrest, we performed flow cytometry-based cell cycle analysis and found a strong G2/M arrest in CH12 but not primary B cells (Fig. 4A,B, Fig. S4A,B). We also measured the expression and phosphorylation of the cell cycle regulator CDC2. Phosphorylation of the cyclin-dependent kinase CDC2 at Tyrosine15 (pTyr15) inhibits kinase activity and serves as a marker of G2/M arrest (58). We observed accumulation of CDC2 and pTyr15-CDC2 in CH12 cells (Fig. 4C), but not in primary B cells (Fig. 4E,G). When normalized to total CDC2 levels, pTyr15-CDC2 showed an increase following PDS exposure only in CH12 cells (Fig 4D,F,H). These results suggest that PDS triggers G2/M arrest in CH12 but not in primary B cells. To further investigate whether cells arrest in G2, we quantified the fraction of cells in mitosis by immunostaining for phosphorylated histone H3 at serine 10 and found no significant difference in the mitotic index of primary cells ((59), Fig 4I,J). Thus PDS exposure also does not impact the mitotic index of primary B cells (Fig. 4J).

**Figure 4.**
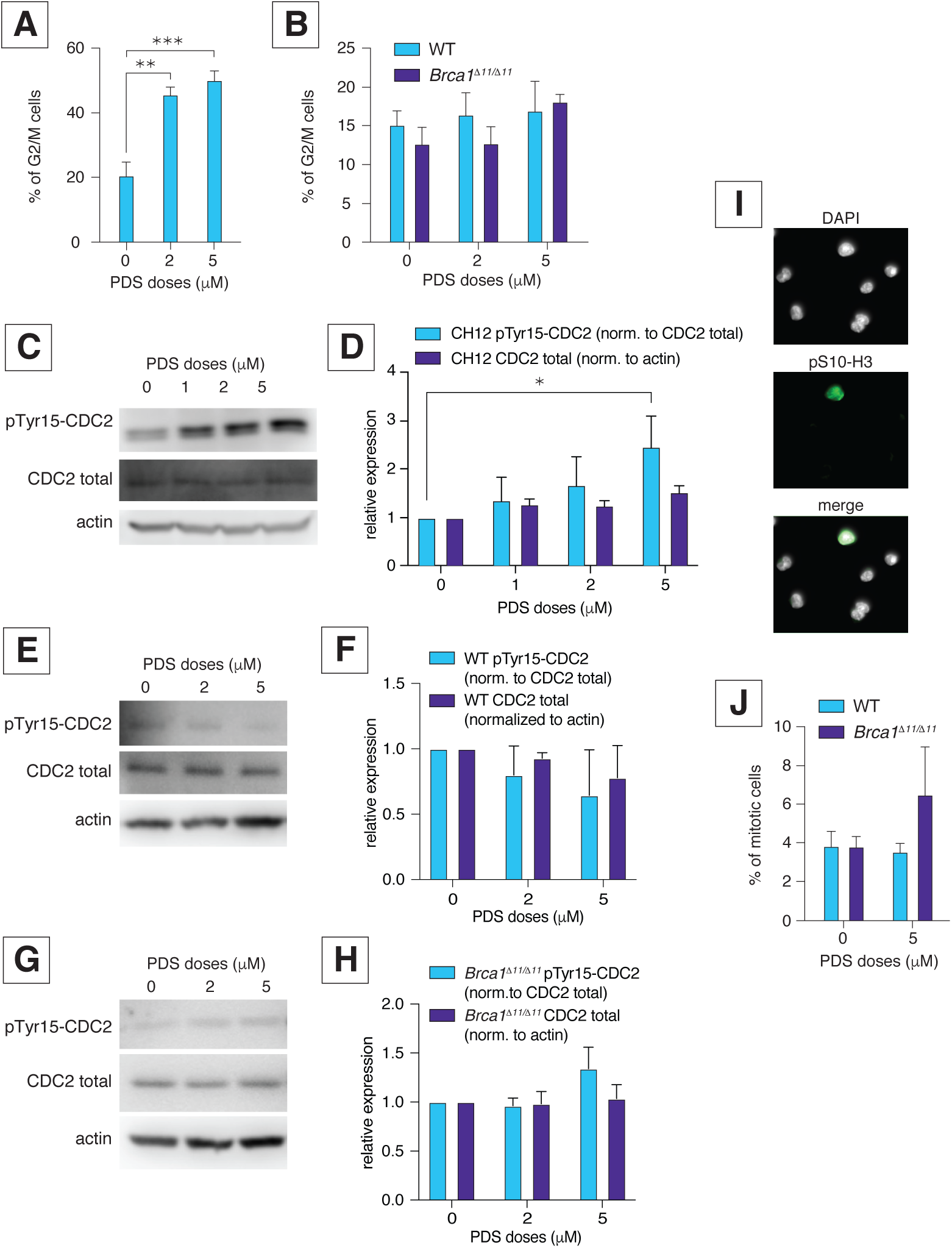
PDS-induced tetraploid metaphases are associated with absence of G2/M arrest. **A)** Percent of CH12 cells in G2/M phase. Error bars show the SEM from three independent experiments. Statistics: **p < 0.01 comparing PDS doses. **B)** Percent of WT and *Brca1^111/111^* primary B cells in G2/M phase. Error bars show the SEM from three independent experiments. **C)** Protein blot analysis of pTyr15-CDC2 and total CDC2 expression in response to PDS in CH12 cells. Actin in each sample was employed as a standard. **D)** Quantification of pTyr15-CDC2 (relative to total CDC2) and total CDC2 (relative to actin) expression in response to PDS in CH12 cells. Error bars show the SEM from three independent experiments. Statistics: *p < 0.05 comparing PDS doses. **E)** Protein blot analysis of pTyr15-CDC2 and total CDC2 expression in response to PDS in WT primary B cells. Actin in each sample was employed as a standard. **F)** Quantification of pTyr15-CDC2 and total CDC2 expression in response to PDS in WT primary B cells. Error bars show the SEM from three independent experiments. **G)** Protein blot analysis of pTyr15-CDC2 and total CDC2 expression in response to PDS in *Brca1^111/111^* primary B cells. Actin in each sample was employed as a standard. **H)** Quantification of pTyr15-CDC2 and total CDC2 expression in response to PDS in *Brca1^111/111^* primary B cells. Error bars show the SEM from three independent experiments. **I)** Representative images of H3 pSer10 immunostaining of WT cells in response to 5 µM PDS. H3 pSer10 visualized in green, DAPI is in greyscale. **J)** Mitotic index of WT and *Brca1^111/111^* cells in response to PDS. Error bars show the SEM from three independent experiments.

To determine if a lack of G2/M arrest and mitotic slippage drives PDS-induced tetraploidization in primary B cells, we exposed primary B cells to both PDS and the CDK1 inhibitor RO-3306. CDK1 inhibition arrests cells in late G2 and prevents entry into mitosis (60), creating an artificial G2/M arrest. We exposed cells to PDS in the presence of RO-3306 for 47 hours, then released cells from RO-3306 for 1 hour in the presence of PDS and colcemid (Fig.5A, left). We did not find any tetraploid metaphases in response to combination of PDS and RO-3306 (Fig. 5A), highlighting a critical role for G2/M arrest in preventing PDS-induced tetraploidy. Treatment with PDS+RO-3306 did not significantly increase chromosomal instability compared to PDS alone (Fig. S5A). We also found a dramatic decrease in the dicentric chromosome frequency in response to PDS+RO-3306 (Fig. 5B-C). These results indicate that majority of dicentric chromosomes also form over 2 or more cell cycles.

**Figure 5.**
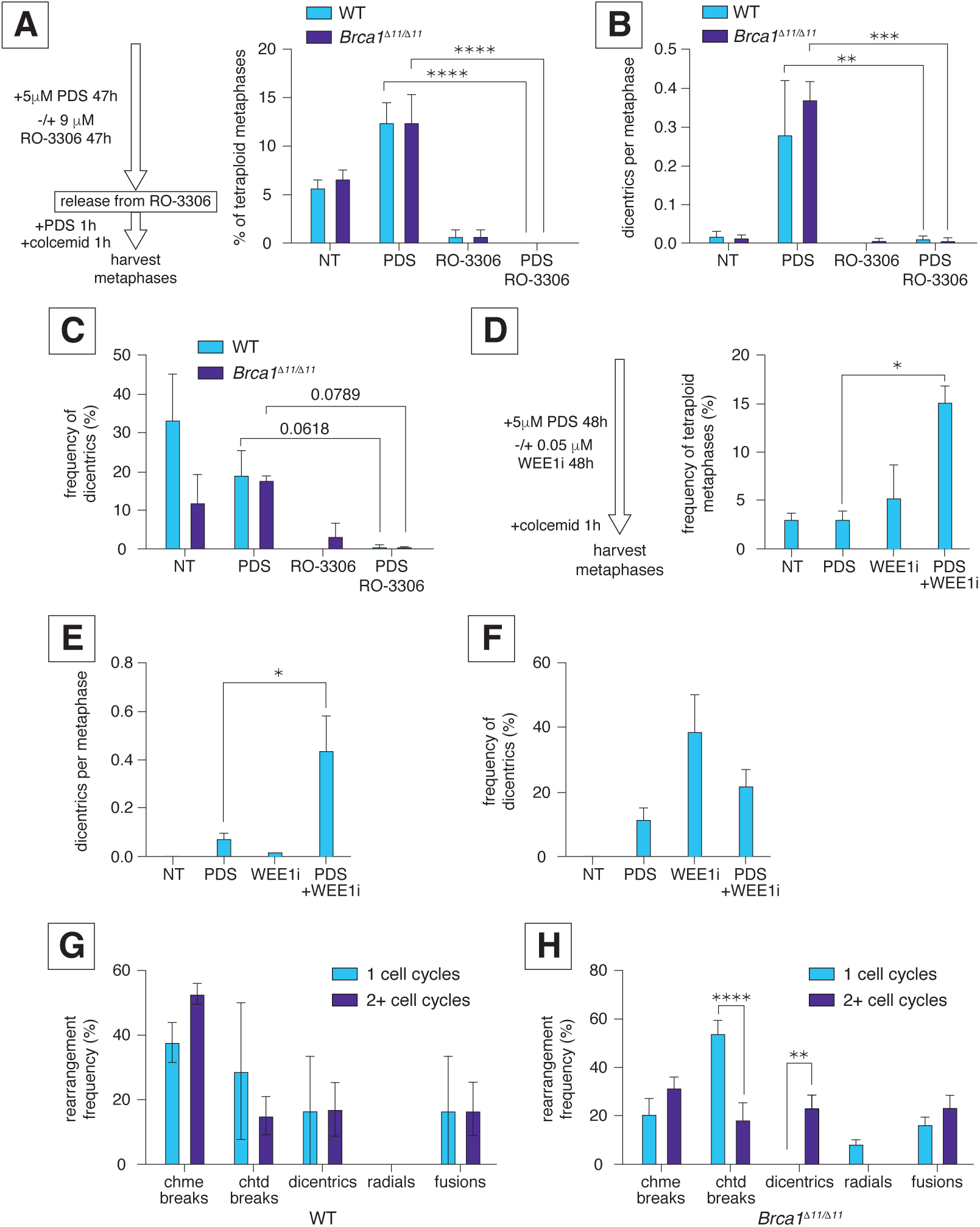
PDS treatment during a single cell cycle does not cause tetraploidy or extensive dicentric chromosome formation. **A)** Left pane – scheme of the experiment. Right pane - percent of tetraploid metaphases in response to PDS combined with RO-3306 in WT and *Brca1^111/111^* cells. Error bars show SEM from three independent experiments. Statistics: ****p < 0.001 comparing 5 µM PDS and combination of 9 µM RO-3306 with 5 µM PDS. **B)** Number of dicentric chromosomes per metaphase in response to PDS combined with RO-3306 in WT and *Brca1^111/111^* cells. Error bars show the SEM from three independent experiments. Statistics: ***p < 0.005 and **p < 0.01 comparing 5 µM PDS and combination of 9 µM RO-3306 with 5 µM PDS. **C)** Dicentric chromosome frequency (%) in response to PDS combined with RO-3306 in WT and *Brca1^111/111^* cells. Error bars show the SEM from three independent experiments. Statistics: exact p-values are shown comparing 5 µM PDS and combination of 9 µM RO-3306 with 5 µM PDS. **D)** Left pane – scheme of the experiment. Right pane - tetraploid metaphase frequency (%) in response to PDS combined with AZD1775 in CH12 cells. Error bars show the SEM from three independent experiments. Statistics: *p < 0.05 comparing 5 µM PDS and combination of 0.05 µM AZD1775 with 5 µM PDS. **E)** Number of dicentric chromosomes per metaphase in response to PDS combined with AZD1775 in CH12 cells. Error bars show the SEM from three independent experiments. Statistics: *p < 0.05 comparing 5 µM PDS and combination of 0.05 µM AZD1775 with 5 µM PDS. **F)** Dicentric chromosome frequency (%) in response to PDS combined with AZD1775 in CH12 cells. Error bars show the SEM from three independent experiments. **G)** Frequencies of different PDS-induced chromosomal rearrangements during only one cell cycle and few cell cycles in WT B cells exposed to 5 µM PDS. Error bars show the SEM from three independent experiments. **H)** Frequencies of different PDS-induced chromosomal rearrangements during only one cell cycle and few cell cycles in *Brca1^111/111^*B cells exposed to 5 µM PDS. Error bars show the SEM from three independent experiments. Statistics: ****p < 0.001 and **p < 0.01 comparing one cell cycle to two or more cell cycles.

We hypothesized that disrupting G2/M arrest in CH12 cells in response to PDS will allow tetraploid cells to enter metaphase. We inhibited PDS-induced G2/M arrest in CH12 cells using the WEE1 inhibitor AZD1775 (Fig. 5D, left). WEE1 kinase inhibition (WEE1i) prevents inactivation of CDK1 and CDK2, disrupting intra-S and G2/M checkpoint activation thus cells readily enter into mitosis (60). Combined exposure to PDS and WEE1i significantly increased chromosomal instability in CH12 cells compared to PDS alone (Fig. S5B). We also observed significantly more tetraploid metaphases (Fig. 5D) and dicentrics in response to PDS+WEE1i than PDS alone (Fig. 5E-F). These results further support our hypothesis that PDS-induced tetraploidy in mouse primary B cells arises from the absence of G2/M arrest.

### PDS treatment during a single cell cycle does not cause tetraploidy or extensive dicentric chromosome formation

Exposure to PDS for 48 hours affects 2 or more cell cycles in B cells, therefore chromosome rearrangements could arise from DNA damage and repair across multiple cell cycles. CDK1i treatment arrested B cells in G2/M, restricting cells to a single cell cycle in the presence of PDS. This arrest caused the absence of tetraploidization and suppressed dicentric chromosome formation (Fig. 5A-C). To characterize chromosomal instability during a single cell cycle, we treated freshly-isolated naïve G0 B cells with PDS for 26 hours in the presence of BrdU to label metaphases that underwent only one round of replication (61) (Fig. S5C; equal labeling of sister arms). Again, *Brca1^111/111^* cells exhibited much higher instability than WT cells (Fig. S5D). We found no tetraploid metaphases. Interestingly, PDS-induced rearrangements in one cell cycle were different compared to 2+ cell cycles; the difference was particularly significant in *Brca1^111/111^* cells due to the higher rate of aberrations (Fig. S5D-E). After one cell cycle in PDS, unrepaired breaks mostly localized to pericentromeric regions, and we observed significantly more chromatid breaks than cells experiencing 2+ cell cycles (Fig. 5G-H). Dicentrics frequency was significantly lower in *Brca1^111/111^* cells after one cell cycle compared to 2+ cell cycles (Fig. 5G-H). We suggest that PDS primarily causes pericentromeric chromatid breaks in the first cell cycle that in subsequent cell cycles turn into chromosome breaks and later to 2-arm fusions including dicentric chromosomes (Fig. 7H).

### PDS causes pericentromeric breakage in immortalized and transformed human B cell lines

Drug targeting specific G4 regions in B cell lymphomas can suppress tumor growth (62, 63), therefore we investigated if PDS induces instability in human B cells using EBV-immortalized GM12878 cells, Raji Burkitt’s lymphoma cells, and PMBCs from healthy donors. We found that PDS exposure induces apoptosis and chromosomal instability in a dose-dependent manner in GM12878 and Raji cells, while in PMBCs it did not cause chromosomal instability even at very high doses (Fig. 6A, Fig. S6A,B). Interestingly, high PDS doses (20 µM and higher) caused erythrocyte coagulation of peripheral blood samples, while PMBCs proliferated normally. Similar to observations in mouse B cells, we found frequent pericentromeric breaks (∼40%) in GM12878 and Raji cells (Fig. 6B-C). This observation is consistent with previous studies showing that G4 structures impair replication fork progression in human pericentromeres (64). Interestingly, we found PDS-induced tetraploid metaphases only in GM12878 cells, but not in Raji cells or PMBCs (Fig. 6D). We also observed a significant increase in dicentrics in response to PDS in GM12878 cells, while in Raji cells the frequency of PDS-induced dicentrics did not significantly differ from spontaneous levels (Fig. 6E-F). The increase in the total number of dicentrics in PDS-treated Raji cells may be non-specific and can be explained by very high rate of PDS-induced total chromosomal instability (Fig. 6E-F, Fig. 6A). We conclude that PDS affects immortalized and transformed human B cell lines causing chromosomal instability, making it a potential therapeutic agent for B cell cancers.

**Figure 6.**
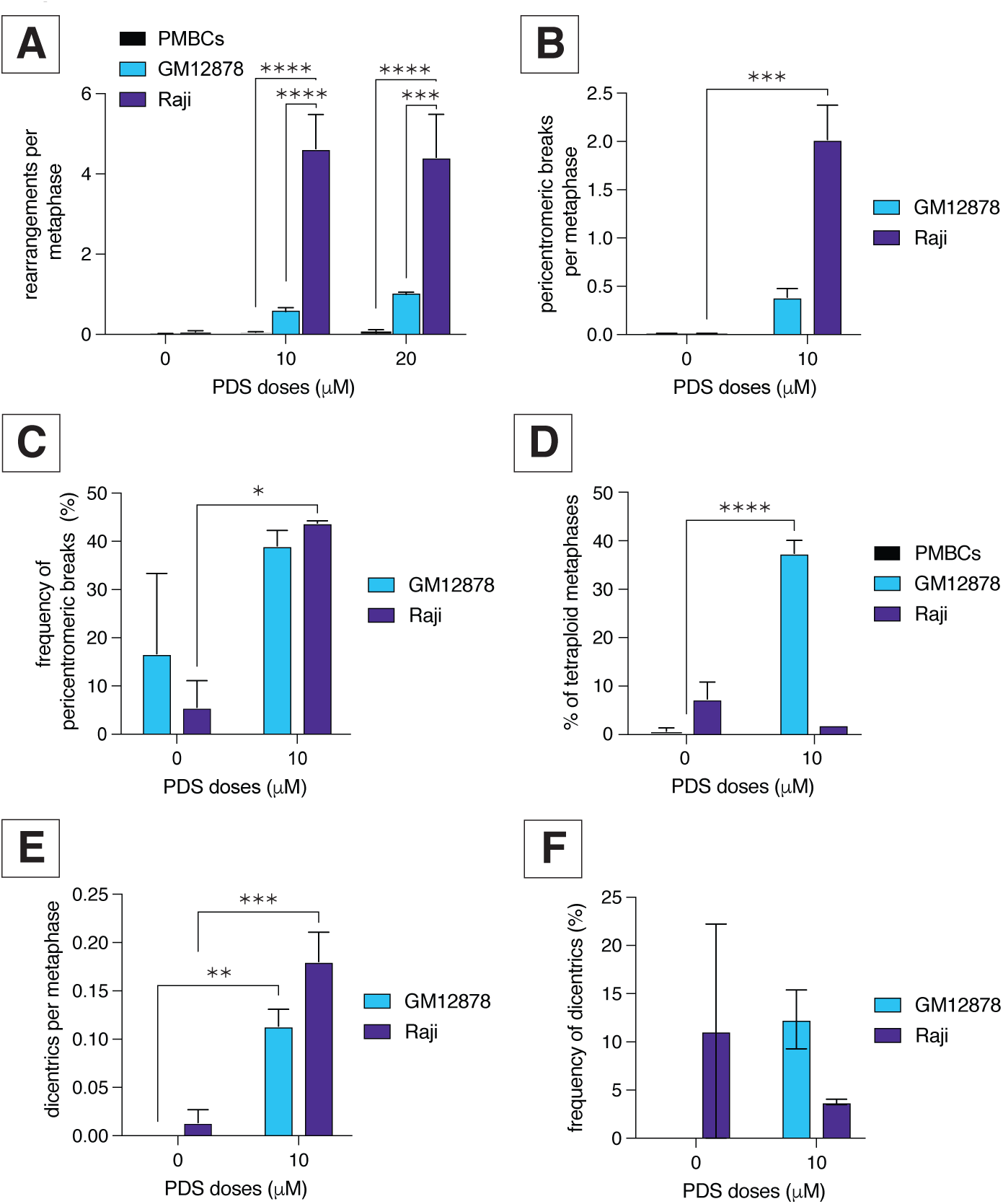
PDS causes pericentromeric breakage in human B cell lymphoma cell lines. **A)** Number of DNA aberrations per metaphase in response PDS in PMBCs, GM12878, and Raji cells. Error bars show SEM from three independent experiments. Statistics: ***p < 0.005 and ****p < 0.001 comparing PDS doses. **B)** Pericentromeric breaks per metaphase in response to PDS in GM12878 and Raji cells. Error bars show SEM from three independent experiments. Statistics: ***p < 0.05 comparing PDS doses. **C)** Frequency of pericentromeric breaks in response to PDS in GM12878 cells. Error bars show SEM from three independent experiments. Statistics: *p < 0.05 comparing PDS doses. **D)** Percent of tetraploid metaphases in response to PDS in PMBCs, GM12878, and Raji cells. Error bars show SEM from three independent experiments. Statistics: ****p < 0.001 comparing PDS doses. **E)** Number of dicentric chromosomes per metaphase in response to PDS in GM12878 and Raji cells. Error bars show the SEM from three independent experiments. Statistics: ***p < 0.005 and **p < 0.01 comparing PDS doses. **F)** Dicentric chromosome frequency (%) in response to PDS in GM12878 and Raji cells. Error bars show the SEM from three independent experiments. Dicentric chromosome frequency (%) in response to PDS in Raji cells. Error bars show the SEM from three independent experiments. Statistics: *p < 0.05 comparing PDS doses.

To determine whether PDS-induced pericentromeric breakage is tissue-specific and restricted to B cell lines, we treated human bone osteosarcoma (U2OS) and breast adenocarcinoma (MCF7) cell lines with 5 µM PDS. Both cell lines exhibited a tendency toward PDS-induced pericentromeric fragility and metaphase tetraploidy, although these effects were not statistically significant (Fig. S6. C-H). Notably, both U2OS and MCF7 cell lines display high baseline chromosomal instability, particularly characterized by numerous mini-chromosomes. As a result, the extent of pericentromeric damage observed may be underestimated due to limitations inherent in gross chromosomal analysis; most of the damage we could classify as *de novo* pericentromeric damage were chromatid and chromosome breaks. We conclude that PDS-induced pericentromeric fragility can occur in other tissues, but B cells appear disproportionately affected. Further experiments in other primary cells will help delineate tissue-specific effects of G4 ligand instability.

### Pericentromeric breakage is induced by G4 stabilization

While PDS is relatively selective for G4s (18), the PDS-induced effects we observed may be caused by off-target binding to non-G4 DNA structures (65). To determine if pericentromeric breakage and tetraploidy are specific to G4 stabilization we analyzed effects of another two G4 stabilizers, CX-5461 ((23)Trial ID: ACTRN12613001061729; Canadian Cancer Trials Group ID: NCT02719997) and Phen-DC3 (66) in mouse primary B cells and CH12. CX-5461 induced DNA damage in both WT and *Brca1^111/111^* cells, particularly in pericentromeric regions (Fig. 7A-C). FISH analysis with rDNA and MaSat probes in WT cells shows that ∼35% of aberrations occur near rDNA, and ∼30% in MaSat regions (Fig.7D-F). CX-5461 also induced tetraploidy correlated with dicentric chromosome formation in *Brca1^111/111^* cells, but the increase was not significant (Fig. S7A-C). PhenDC3 also induced DNA damage with a high percentage at pericentromeric regions, but the increase in damage was not significant (Fig. S7D-H). Taken together, these findings demonstrate that pericentromeric fragility is a consequence of G4 stabilization while extensive tetraploidization may be specific to PDS.

**Figure 7.**
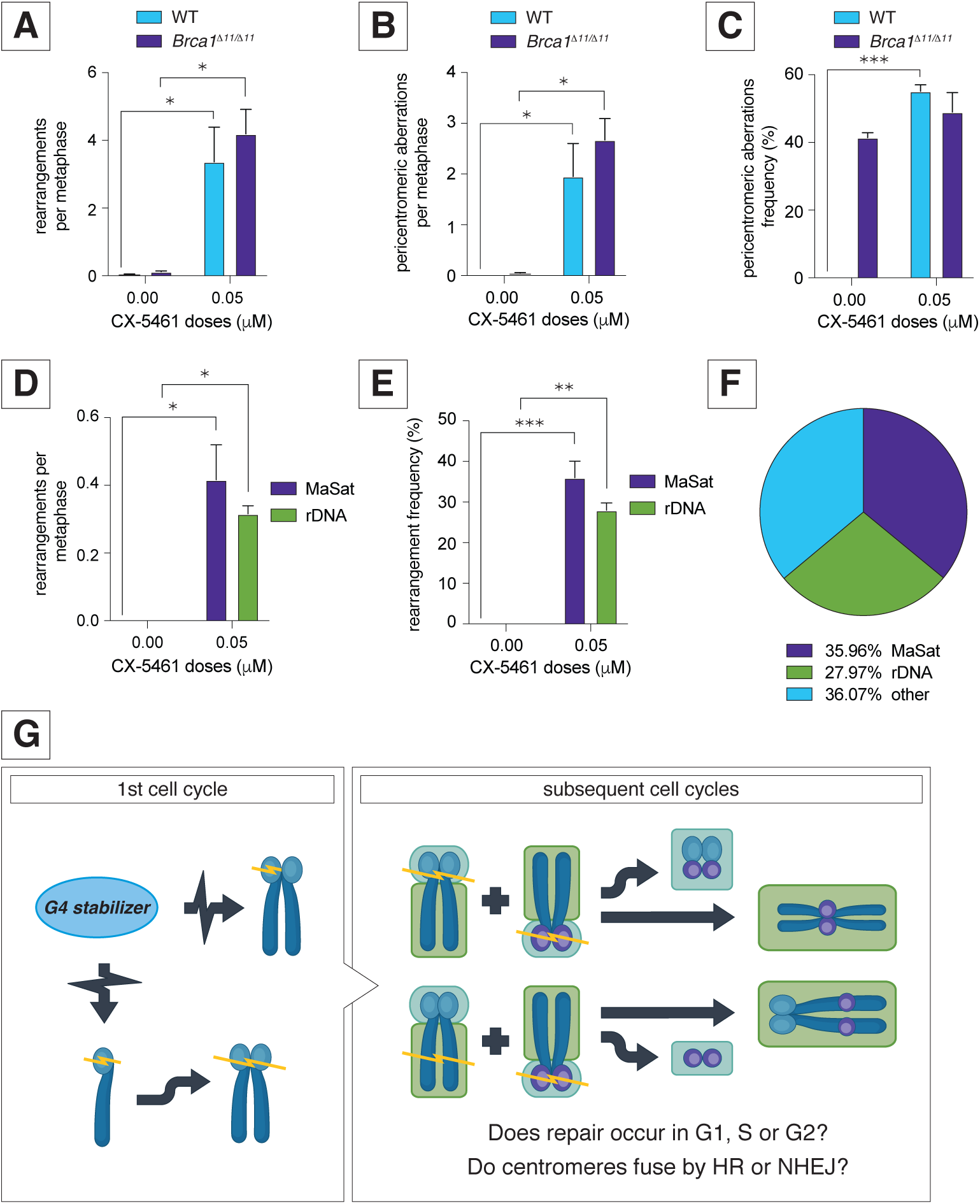
G4 stabilizer CX-5461 induces specific chromosomal rearrangements similar to those caused by PDS in WT, *Brca1^111/111^* cells. A) Number of DNA aberrations per metaphase in response to CX-5461 in WT and *Brca1^111/111^* cells. Error bars show SEM from three independent experiments. Statistics: *p < 0.05 comparing CX-5461 doses. **B)** Pericentromeric breaks per metaphase in response to CX-5461 in WT and *Brca1^111/111^* cells. Error bars show SEM from three independent experiments. Statistics: *p < 0.05 comparing CX-5461 doses. **C)** Frequency of pericentromeric breaks in response to CX-5461 in WT and *Brca1^111/111^*cells. Error bars show SEM from three independent experiments. Statistics: ***p < 0.005 comparing CX-5461 doses. **D)** MaSat and rDNA aberrations per metaphase in response to CX-5461 in WT cells. Error bars show SEM from three independent experiments. Statistics: *p < 0.05 comparing CX-5461 doses. **E)** Frequency of MaSat and rDNA aberrations in response to CX-5461 in WT cells. Error bars show SEM from three independent experiments. Statistics: **p < 0.01 and ***p< 0.005 comparing CX-5461 doses. **F)** Fraction of DNA rearrangements at rDNA and MaSat in response to 0.05 µM CX-5461 in WT cells. **G)** Working model of G4 ligand-mediated genome instability in B cells. G4 stabilization induces complex rearrangements, especially pericentromeric rearrangements, over two or more cell cycles. Chromatid breaks in pericentromeric regions occur in a single cell cycle and are inherited by a daughter cell. In subsequent cell cycles, broken chromatids can either be fused by NHEJ or HR in G1 or replicated and then fused in S/G2. Both G1 and S/G2 fusions can create multicentromeric and/or acentric rearrangements.

## Discussion

G4s are a natural replication barrier and require the orchestrated work of helicases, DNA replication and DNA repair machineries to ensure faithful and error-free replication. Cancer cells frequently harbor mutations in DNA repair factors inducing chronic replication stress. Thus, G4 stabilization in cancer cells can be a significant obstacle for replication leading to chromosomal instability and cell death. Here we identified repetitive DNA elements as genomic loci sensitive to PDS-induced chromosomal instability in WT and BRCA1-deficient mouse B cells. Expanding our studies to mouse CH12 and human B cells, we showed PDS-induced pericentromeric instability is conserved in mouse and human cancers. Taken together, our findings suggest that G4 stabilization can be potential target for B cell cancer treatment.

### G4 stabilization induces damage at pericentromeric and rDNA repeats

PDS exposure induces recurrent damage at two major repeat sequences in mouse cells: rDNA repeats and the MaSat repeats found on all chromosomes. This preponderance of damage at MaSat was not observed in studies examining chromosome fragility and rearrangements from other replication stress-inducing agents including the B-type polymerase inhibitor aphidicolin, the ribonucleotide reductase inhibitor hydroxyurea, or inhibition of the ataxia telangiectasia and Rad3 related (ATR) checkpoint kinase (61, 67, 68). In both mouse and human, the rDNA unit exhibits strong G4-forming capability throughout the 47S (with a high frequency of G4s in throughout the coding region and the 3’ enhancer / promoter region) and intergenic spacer regions (38, 69) (Table S1). PDS was shown to induce R-loop accumulation in cancer cells (31), which potentially can lead to chromosomal instability in such actively transcribed regions like rDNA. Also, rDNA transcription was shown to contribute to the cellular response to G4 stabilization: rDNA transcription suppression completely abrogated PDS-induced DSB formation (70); while active rDNA transcription sensitized cells and increased DNA damage occurrence in response to CX-5461 (71). The rDNA repeats are the most actively transcribed regions in the genome, thus collisions between transcription and replication in stabilized G4-forming areas can lead to accumulation of R-loops, stalled replication forks, double-strand breaks, and further rDNA and total genomic instability.

Secondary structure is a shared feature of eukaryotic centromeric DNA, particularly in metazoans, and is proposed to be a key part of centromere identity (72). Unfortunately, there are no studies about possible G4 structures in mouse MaSat and MiSat DNA. Using G4 Hunter we found that the pericentromeric mouse MaSat harbors a single, potential G4-forming sequence that is relatively weak, with only two planar tetrads and one long loop sequence (Table S1). In humans, G4 and hairpin structures are detectable in the pericentromeric area of chromosome 9 and in pericentromeric satellite 3 sequences (73, 74). Interestingly, Mendez-Bermudez and colleagues showed that the shelterin subunit TRF2 binds to satellite 3 sequences during S phase, allowing the recruitment of the G4-resolving helicase RTEL1 to facilitate fork progression (64). Human pericentromeric satellite II, found in abundance on chromosomes 1 and 16, also has weak G4-forming potential (75) (Table S1). These observations suggest that pericentromeric satDNA is enriched in G4 structures that — when affected by enhanced transcription by demethylation or stabilized by G4 ligands — impede or disrupt replication, leading to chromosomal rearrangements. Of note, we find that PDS can induce damage within a single cell cycle (Figure 5G,H), suggesting that DSB formation does not predominantly occur during chromosome separation in mitosis. In contrast, the under-replication of DNA and subsequent rupture of ultra-fine and anaphase DNA bridges during mitosis, enhanced by exposure to low dose aphidicolin or hydroxyurea, is thought to be a major cause of DSB formation at late-replicating common fragile sites (76–78). Such ruptures would lead to DNA breaks that persist into the second cell cycle that are visible as 53BP1 nuclear bodies, large accumulations of 53BP1 and other DNA repair factors often found in both daughter cells (79, 80). Instead, we propose breaks occur in one cell cycle, and daughter cells inherit chromosomes with breaks in pericentromeric and other regions (Figure 7G). These broken chromosomes fuse in subsequent cell cycles, either by non-homologous end-joining (NHEJ) or homologous recombination (HR). HR can occur in G1 and non-cycling cells in centromeric repeats (81, 82), therefore it will be interesting to determine how the cell cycle impacts repair pathway choice and cell survival.

PDS induced damage in pericentromeric repeats suggests G4-forming elements in these regions, but their detection has largely been missed by next-generation sequencing (NGS) methods (5, 7, 83). This discrepancy can be attributed to limitations of NGS and inherent bias in data analysis – most pipelines discard reads mapping to more than one genomic region. The new telomere-to-telomere map of human chromosomes will help clarify where G4 sequences form, their relative prevalence in the genome and the impact G4s have on the epigenetic and transcriptional state of repetitive regions (29, 75, 84, 85). This breakthrough will help interpret current data analyzing G4 ligand effects under a variety of conditions and tissue/cell types including telomerase/TERRA expression, proliferative rate, and alterations to transcriptional programs in development, differentiation, and stress.

### G4 stabilization induces tetraploidization

Polyploidy is physiological for some cell types, but is associated with tumorigenesis and cancer in normally diploid cells (54, 86, 87). Tetraploidy is one of the most common stages of polyploidy (86). We found that PDS induces tetraploidy in primary and transformed mouse B cells and in immortalized human GM12878 B cells. This is consistent with previous studies showing that failure to remove stabilized G4s in telomeres results in tetraploidy and aneuploidy (88). PDS causes replication stress, which in turn can induce chromosomal instability and tetraploidy (89). The majority of tetraploids produced by mitotic slippage undergo apoptosis in timely manner (90), but some tetraploid cells can escape and contribute to tumorigenesis (86). In mouse CH12 cells, we observed PDS-induced tetraploidy in interphase nuclei, but not in metaphases, which was associated with strong G2/M arrest. We suggest that this G2/M arrest can be first step in the route of apoptosis, though more studies are needed to confirm this hypothesis. The extensive PDS-induced tetraploid metaphases in mouse primary B cells and human GM12878 cell line may be evidence of G2/M checkpoint escape. However tetraploid metaphases did not significantly increase in the Burkitt’s lymphoma Raji cell line. These observations suggest that tetraploidization combined with the ability to enter mitosis to continue proliferation is a frequent event only under specific circumstances; thus, the molecular pathways governing PDS-induced tetraploidization and subsequent proliferation should be investigated further in multiple cell types.

PDS, like most chemotherapeutic agents, is likely a double-edged sword and may initiate tumorigenesis through tetraploidization. In B cell cancers, this may relate to how cells are activated as different stimuli promote cell viability and suppress apoptosis using overlapping and distinct pathways (91–94). It would be interesting to investigate whether different forms of stimulation differentially impact PDS-mediated genome instability and tetraploidization.

### PDS as cancer therapeutic agent

Since G4s are involved in carcinogenesis (4, 95–97), significant effort has been made to develop G4-targeting agents as cancer therapeutics (98). Ligand-mediated G4 stabilization can suppress replication, transcription, and oncogenes expression in malignant cells (98). To date, one G4 ligand in ongoing human clinical trials, CX-5461, has been reported to successfully treat blood cancers and BRCA1/2-deficient tumors (23, 99, 100).

We suggest PDS and its derivatives are also potential cancer therapeutics. PDS causes DNA damage and apoptosis in BRCA1/2-deficient cancer cells, including cells resistant to PARP inhibition (22). PDS has also been reported to reduce SRC proto-oncogene expression (19). Recent genome-wide shRNA screen for PDS sensitivity identified genes related to cell cycle, ribosome, and DNA replication (101). These findings are consistent with our results showing that PDS can cause chromosomal instability, including rDNA damage, as well as G2/M arrest in rapidly proliferating cell types. Cancer cells frequently exhibit increased rDNA transcription (102), suggesting that PDS-mediated transcription issues and genomic instability of rDNA may effectively decrease malignant cells viability and proliferation.

Hematological cancers often show fragility in pericentromeric satDNA, a hotspot of translocations and gross chromosomal rearrangements (103, 104). According to our study, PDS largely affects pericentromeric satDNA; thus PDS treatment of cancer cells may exacerbate already pre-existing satDNA vulnerability and cause extensive satDNA breakage leading to further cell death. Another attractive strategy is enhancing G4 ligand sensitivity of DSB repair-defective cancers by combining with existing therapies that also cause genome instability. Small molecule inhibition of WRN sensitizes cells to the G4 ligand telomestatin (105). Suppression of non-homologous end-joining by DNA-PKcs inhibition also dramatically sensitizes cells to both PDS and CX-5461 (20, 23). Here, combined PDS and WEE1 inhibitor exposure dramatically increases chromosomal instability compared to PDS alone in CH12 lymphoma cells, suggesting potential synergy of G4 ligands with drugs targeting cell cycle checkpoints.

Though PDS is considered a promising anti-cancer agent, possible side effects and drug-induced cancer require attention. We show that PDS does not cause cytoxicity or chromosomal instability in human PBMCs, indicating it is a potentially safe candidate for anti-cancer therapy. Unfortunately, Moruno-Machon and colleagues showed PDS cytotoxicity in primary neuronal cells, however it is unknown if neurological side effects will be observed *in vivo* (32). Another concern is PDS-induced tetraploidization, which can be a cancer-initiating event. Taken together, G4 stabilization by PDS is promising target for blood cancers treatment but further *in vivo* studies are required for successful implementation.

## Materials and Methods

### Mice and cells

All experiments were performed in accordance with protocols approved by the UC Davis Institutional Animal Care and Use Committee (IACUC protocol #23444). Mice used in this study include CD19-cre, and *Brca1^111/111^* (34, 33). Splenic B cells were isolated and cultured as previously described (68). CH12, GM12878, and Raji cells were cultured according to standard protocols. Human PBMCs were obtained from three unrelated volunteers and cultured for 72 h according to standard protocols. Additional details can be found in SI.

### Chemical compounds

The experiments were performed using PDS (Pyridostatin Hydrochloride, Millipore Sigma, SML2690), CX-5461 (Millipore Sigma, 509265), Phen-DC3 (Millipore Sigma, SML2298), BrdU (Sigma, B5002), RO-3306 (Millipore Sigma, SML0569), and AZD1775 (Adavosertib, MedChemExpress, HY-10993).

### FISH Probes and BrdU-detection

All Bacterial Artificial Chromosomes (BACs) used for custom-designed probes were purchased from BACPAC Genomics: RP23-225M6 (rDNA) (106) and RP23-405G16 (BCL2) (61). The MaSat plasmid (a 471 bp fragment of MaSat (107) cloned into pBluescript II KS) was a gift by Professor Olga I. Podgornaya (Saint-Petersburg, Russia). Telomere probe (Tel-Cy3, F1002) and centromere probe (CENPB-Alexa488, F3004) colocalizing with MiSat were purchased from PNAbio. For BrdU detection, the primary mouse-anti-BrdU (BD, 347580; 1:200) and secondary Cy5 goat-anti-mouse antibodies (Invitrogen, A10524; 1:200) were used.

### Immunofluorescence

Primary histone H3 [p Ser10] antibody (Novus Biologicals, NB21-1091; 1:200) and secondary goat-anti-Mouse IgG (H+L) cross-adsorbed secondary antibody, Alexa Fluor™ 488 (ThermoFisher Scientific, A-11001; 1:200) were employed.

### Immunoblotting

Anti-phospho-CDC2 (Tyr15) (Cell Signaling Technology, 4539; 1:1000), anti-CDC2 total (Cell Signaling Technology, 28439; 1:1000), anti-β-Actin (ABclonal, AC026; 1:5000), and secondary HRP-conjugated mouse anti-Rabbit (ABclonal, AS061; 1:5000) antibodies were employed.

### Statistics

A minimum of 50 metaphases in FISH studies and at least 100 nuclei in TUNEL assay were analyzed in triplicates for each condition. Statistical significance of differences was estimated by Student’s t-criterion or two-way ANOVA. To estimate the correlation between co-occurrence of dicentric chromosomes and tetraploidy in individual cells, the Cochran-Mantel-Haenszel test was used.

Further details can be found in the Supplementary Information.

## Supporting information

Combined supplemental figures and tables

## Acknowledgments

We would like to thank Frederic Chedin and members of the Chedin lab for helpful conversations. Special thanks to Professor Olga I. Podgornaya for the MaSat plasmid and Jack McTiernan for assistance in figure design and layout.

This project was supported by a University of California Cancer Research Coordinating Committee (CRCC) Pilot grant (C23CR5562) and NIGMS R01 (GM134537) to Dr. Jacqueline Barlow. It was also supported by the University of California Davis Flow Cytometry Shared Resource Laboratory with funding from the NCI P30 CA093373 (Comprehensive Cancer Center), and technical assistance from Bridget McLaughlin, Jonathan Van Dyke and Ashley Karajeh.

