## Supplementary material for "G-quadruplex stabilization induces DNA breaks in pericentromeric repetitive DNA sequences in B lymphocytes": Combined supplemental figures and tables

*corresponding author: Jacqueline H. Barlow

**This PDF file includes:**

Supporting text: extended materials and methods
Figures S1 to S7
Legends for Figures S1 to S7
Tables S1 to S2
SI References

#### Supporting Information Text

##### Supporting materials and Methods

**Subhead.** Type or paste text here. You may break this section up into subheads as needed (e.g., one section on “Materials” and one on “Methods”).

##### Mice and cells

All experiments were performed in accordance with protocols approved by the UC Davis Institutional Animal Care and Use Committee (IACUC protocol #23444). Mice used in this study include CD19-cre and *Brca1*$^{\Delta 11/\Delta 11}$ (1–3). Splenic B cells were isolated using the Dynabeads untouched CD43 mouse B cell isolation kit (Thermo Fisher, 11422D) and cultured as previously described (4). Briefly, isolated B cells were cultured in B cell media (BCM: RPMI-1640 supplemented with 10% FBS, 1% L-glutamine, 50 IU/ml penicillin/streptomycin, 1% sodium pyruvate, 0.0004% 2-mercaptoethanol, 10 mM HEPES). B cells were stimulated with lipopolysaccharide (LPS), α-RP105, and interleukin 4 (IL-4) to induce the robust proliferation observed when cells undergo CSR to IgG1. CH12 (subclone F3) (5) B lymphocytes were cultured in a modified RPMI 1640 medium with 4 mM L-glutamine and 20 mM HEPES supplemented with 10% FBS, 100 U/ml penicillin-streptomycin solution, 5% NCTC 109, and 0.0004% 2-mercaptoethanol. Human lymphocytes were obtained from peripheral blood of three unrelated volunteers and cultured for 72 h in MF-Chang medium (Irvine Scientific, 91005), supplemented with 10% heat inactivated FBS and 3 μg/mL phytohemagglutinin (Remel, Inc., 30852701). GM12878 human B-lymphoblastoid cells were cultured in a modified RPMI 1640 medium with 2 mM L-glutamine supplemented with 10% FBS, and 100 U/ml penicillin-streptomycin solution. The Raji Burkitt lymphoma cells were grown in RPMI-1640 with 10% FBS and 100 U/ml penicillin-streptomycin solution. The U2OS human bone osteosarcoma and MCF7 human breast adenocarcinoma cell lines were grown in DMEM supplemented with 10% FBS and 100 U/ml penicillin-streptomycin solution. All cells were kept at 37°C and 5% $CO_2$.

##### Chemical compounds and drug treatments

For most experiments, PDS (Pyridostatin Hydrochloride, Millipore Sigma, SML2690) was added to medium 48 h prior to harvest at the designated concentration. For analysis of PDS treatment for one cell cycle, 5 μM PDS was added to freshly-isolated stimulated naïve B cells for 26 h. To calculate the proliferation rate during one cell cycle, 1 μM BrdU (Sigma, B5002) was added to medium for 26 h -/+ PDS. To induce G2 arrest, 5 μM PDS +/- 9 μM RO-3306 (Millipore Sigma, SML0569) were added to B cells for 48 h. To disrupt G2 arrest, 5 μM PDS +/- 0.05 μM AZD1775 (Adavosertib, MedChemExpress, HY-10993) were added to CH12 cells for 48 h. For the experiments using CX-5461 and Phen-DC3, CX-5461 (Millipore Sigma, 509265 and Phen-DC3 (Millipore Sigma, SML2298) were added to medium 48 h prior to harvest at the designated concentration.

##### Metaphase chromosome preparation

Cells were arrested in metaphase by 1h treatment with 0.1 μg/ml demecolcine (Sigma, D1925), treated with 0.075M KCl, fixed in methanol:acetic acid (3:1), spread onto glass slides, and air-dried.

##### FISH Probes

All Bacterial Artificial Chromosomes (BACs) used for custom-designed probes were purchased from BACPAC Genomics: RP23-225M6 (rDNA) (6) and RP23-405G16 (BCL2). The MaSat plasmid (a 471 bp fragment of MaSat (7) cloned into pBluescript II KS) was a gift by Professor Olga I. Podgornaya (Saint-Petersburg, Russia). Probes were direct-labeled using a nick translation kit (Abbott Molecular, Inc., 07J00-001) with DY-495-dUTP (Dyomics, 495-34) or DY-590-dUTP (Dyomics, 590-34) and hybridized to metaphase cells of a karyotypically normal donor to confirm correct mapping. Telomere probe (Tel-Cy3, F1002) and centromere probe (CENPB-Alexa488, F3004) colocalizing with MiSat were purchased from PNAbio.

**FISH and BrdU-detection**
FISH studies were performed as previously described (8). MaSat and rDNA probes were co-denatured with cells at 75°C for 3 min and incubated overnight at 37°C in a humid chamber. Telomeric and CENPB probes were co-denatured for 3 min with cells at 75°C then incubated for 1 h at 37°C in a humid chamber. For BrdU detection, the primary mouse-anti-BrdU (BD, 347580; 1:200) and secondary Cy5 goat-anti-mouse antibodies (Invitrogen, A10524; 1:200) were used.

**Immunofluorescence**
PDS-treated cells were washed with 1xPBS three times for 5 min, fixed in 4% paraformaldehyde for 12 min, permeabilized with 0.5% Triton X-100 in 1x PBS for 5 min, dropped onto glass slides and air-dried. Cells were washed with 1xPBS for 5 min and incubated with primary histone H3 [p Ser10] antibody (Novus Biologicals, NB21-1091; 1:200) for 1 h in a humid chamber at 37°C. After three 5 min washes in 1x PBS, cells were incubated for 45 min with secondary goat-anti-Mouse IgG (H+L) cross-adsorbed secondary antibody, Alexa Fluor™ 488 (ThermoFisher Scientific, A-11001; 1:200) in a humid chamber at 37°C. The cells were washed three times with 1x PBS then mounted in Vectashield antifade, containing DAPI (Vector Laboratories Inc., H-1200).

**TUNEL assay**
TUNEL was performed using the In Situ Cell Death Detection Kit according to the manufacturer protocol (Roche, 11684795910). At least 100 nuclei per experiment were analyzed by microscopy. At least three independent experiments were performed for each data set.

**Microscopy**
A minimum of 50 metaphases were analyzed for each condition. CH12 cells and Raji cells carry stable Robertsonian rearrangements in each cell, therefore we removed these stable rearrangements from analyses and reported only *de novo* aberrations caused by PDS. Metaphases images were acquired using an epifluorescent Axioimager Z2 microscope with Metafer software (Metasystems Group Inc). Interphase cells were acquired using an epifluorescent Nikon microscope with NIS Elements AR4.40.00 software (Nikon). Downstream analysis used ImageJ32 software (NIH).

**Flow cytometry**
Viability assays were performed as previously described (8). Cell proliferation was measured using Carboxyfluorescein succinimidyl ester (CFSE) staining and cell cycle analysis were performed as described in (9). Briefly, freshly isolated primary B cells were washed and resuspended in 0.1% BSA/PBS at $1 \times 10^7$ cells/ml and labeled with CFSE at a final concentration of 5 μM for 10 min at 37°C. CFSE was quenched with ice-cold RPMI 1640 medium containing 10% FBS and washed twice with BCM. Labeled cells were then cultured in BCM and appropriate stimuli for the times indicated prior to analysis by flow cytometry. Fluorescence activated cell sorting (FACS) analysis was carried out on a Becton Dickinson Canto II flow cytometer (BD Biosciences). Up to 20,000 live cells were analyzed for each condition, and data analysis was performed using FlowJo 10.4 software.

**Immunoblotting**
Protein expression was analyzed 48 h after PDS treatment. Protein lysates were prepared as described in (10) and resolved by SDS-PAGE. Gels were transferred to polyvinylidene difluoride membranes (Genesee Scientific, 83-646R). Intercept blocking buffer (LI-COR, 927-60001) was used for membrane blocking and antibody dilution. Anti-phospho-CDC2 (Tyr15) (Cell Signaling Technology, 4539; 1:1000), anti-CDC2 total (Cell Signaling Technology, 28439; 1:1000), anti-β-Actin (ABclonal, AC026; 1:5000), and secondary HRP-conjugated mouse anti-Rabbit (ABclonal, AS061; 1:5000) antibodies were employed. Band signals were scanned using Amersham Imager 600 (GE Healthcare) after incubation with SuperSignal™ West Pico PLUS Chemiluminescent Substrate (ThermoFisher Scientific, #34577). Band density analysis was performed using ImageJ32 software (NIH).

**Statistics**

Statistical significance of differences was estimated by Student’s t-criterion or two-way ANOVA. To estimate the correlation between co-occurrence of dicentric chromosomes and tetraploidy in individual cells, the Cochran-Mantel-Haenszel test was used.

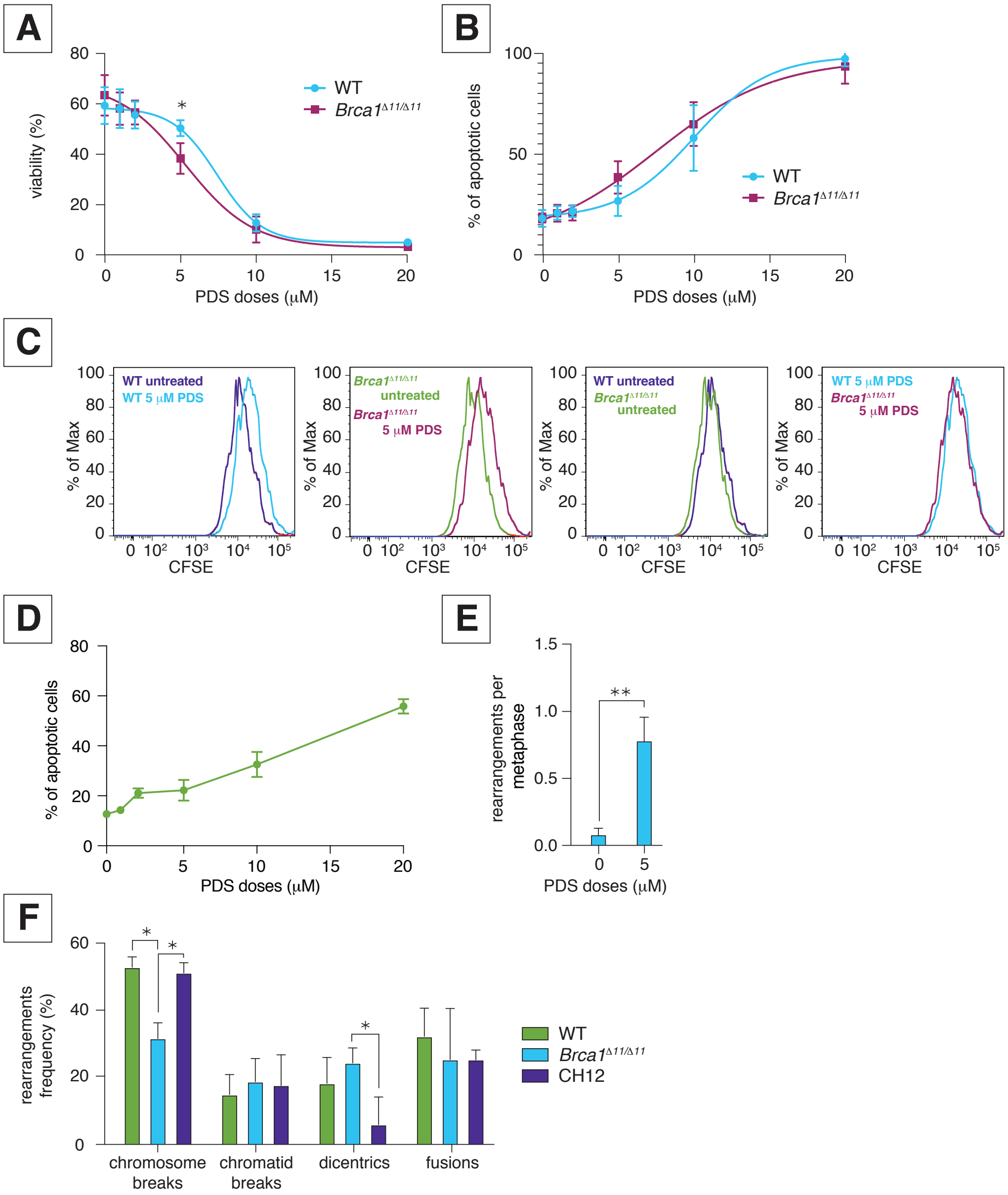

Fig. S1. **WT, *Brca*$^{\Delta 11/\Delta 11}$, and CH12 cells exhibit elevated cell death and chromosomal instability in response to PDS. A)** Viability of WT and *Brca1*$^{\Delta 11/\Delta 11}$ cells in response to PDS treatment measuring cell fragmentation by flow cytometry. **B)** Percent of TUNEL-positive apoptotic WT and *Brca1*$^{\Delta 11/\Delta 11}$ cells in response to PDS treatment. Error bars show the SEM from three independent experiments. *p < 0.05 comparing WT and *Brca1* $^{\Delta 11/\Delta 11}$ cells exposed to PDS. **C)** CFSE histograms of untreated and 5 μM PDS treated WT and *Brca1*$^{\Delta 11/\Delta 11}$ cells. **D)** Percent of TUNEL-positive apoptotic CH12 cells in response to PDS treatment. Error bars show the SEM from three independent experiments. **E)** Number of DNA aberrations per metaphase in response to PDS in CH12 cells. Error bars show SEM from three independent experiments. Statistics: **p < 0.01 comparing PDS doses. **F)** Frequency of specific rearrangements in response to PDS in WT, *Brca1*$^{\Delta 11/\Delta 11}$, and CH12 cells. Error bars show the SEM from three independent experiments. *p < 0.05 comparing WT, *Brca1*$^{\Delta 11/\Delta 11}$, and CH12 cells exposed to PDS.

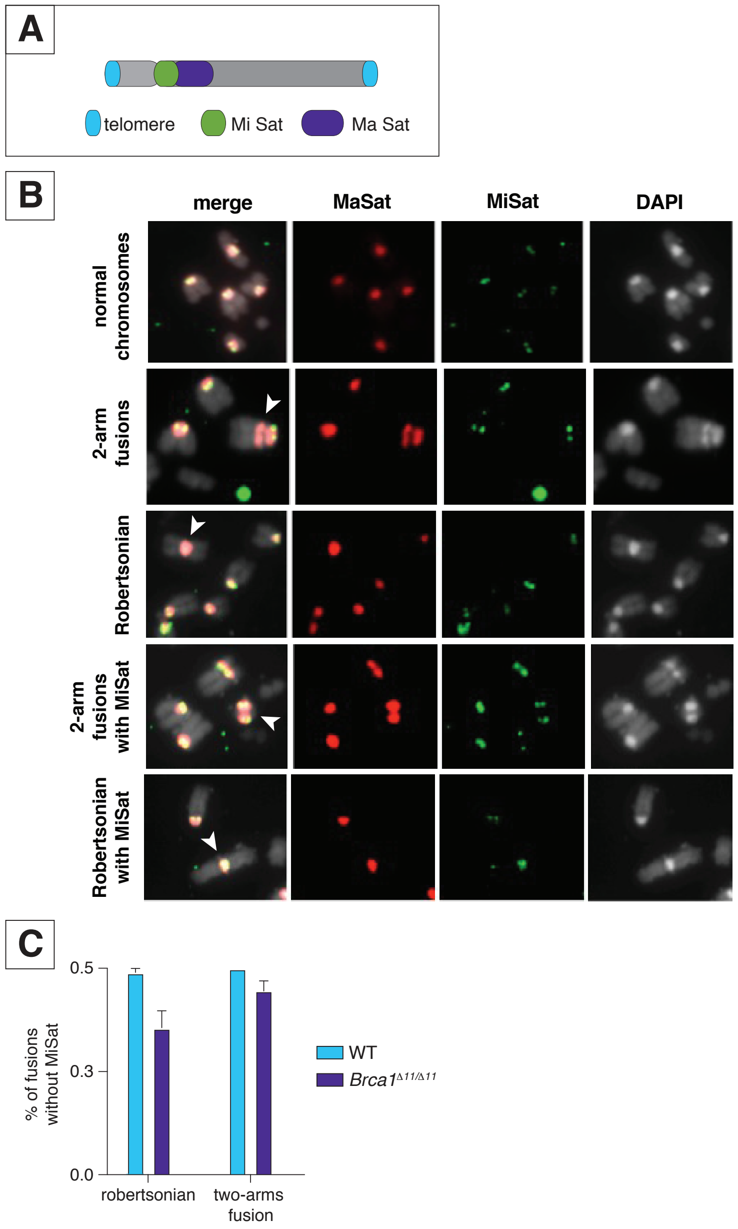

Fig. S2. **SatDNA is involved to formation of Robertsonian chromosomes and two-arms fusions in response to PDS. A)** Scheme of localization of SatDNA repeats on mouse chromosomes. **B)** Representative images of Robertsonian chromosomes and two-arms fusions in response to PDS. Arrows highlight the rearrangements. MaSat probe visualized in red, MiSat is in green, DAPI is in greyscale. **C)** Percent of pericentromeric fusions involving MaSat, but not MiSat, in response to 5 μM PDS in WT and *Brca1*$^{\Delta 11/\Delta 11}$ cells. Error bars show the SEM from three independent experiments.

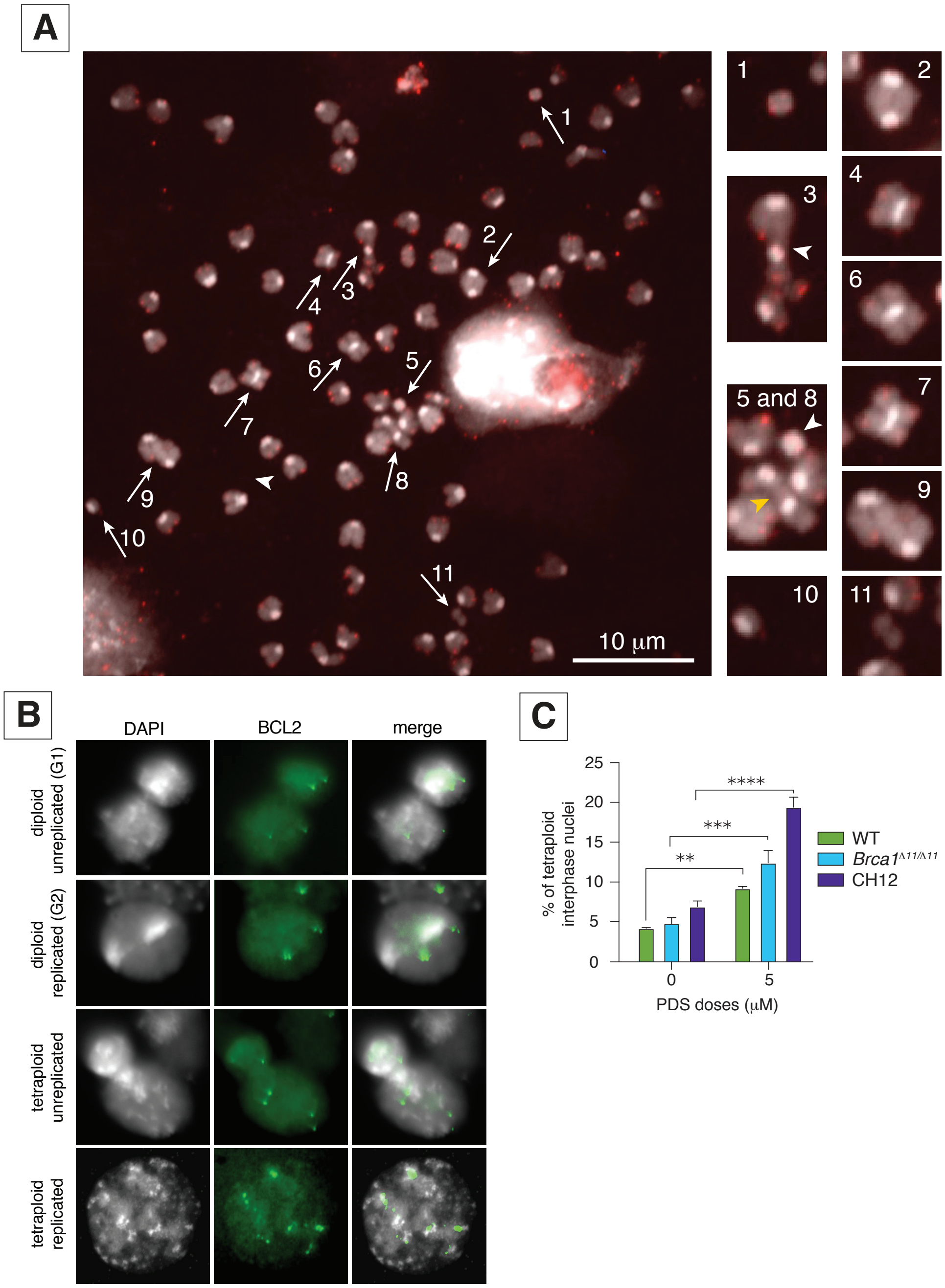

Fig. S3. **CH12 cells show high levels of tetraploid interphase nuclei in response to PDS. A)** Representative image of tetraploid metaphase spread with many of the typical chromosomal rearrangements observed in response to PDS. Numbered rearrangements are enlarged and shown individually on right panels. Arrows highlight the rearrangements: #1 – tiny dicentric chromosome, formed of two centromeres resulted from pericentromeric chromosome breaks; #2 – dicentric chromosome; #3 – pericentromeric chromatid break, followed by fusion with another chromosome's arm; #4, #6, and #7 – Robertsonian chromosomes; #5 – pericentromeric chromosome break (white arrowhead on zoomed in panel); #8 and #9 –1-arm fusions (#8 is yellow arrowhead on right zoomed in panel); #10 and #11 – non-pericentromeric chromosome breaks. Telomere-specific probe visualized in red, DAPI is in greyscale. Arrows highlight the rearrangements; blue arrow highlights small heterochromatic dicentric chromosome. Telomere-specific probe visualized in red, DAPI is in greyscale. **B)** Representative pictures of interphase diploid unreplicated (G1, top panel), diploid replicated (G2, the second panel from the top), tetraploid unreplicated (the third panel from the top), and tetraploid replicated (the bottom panel) B cell nuclei using site-specific BCL2 FISH probe. BCL2 probe visualized in green, DAPI is in greyscale. **C)** Percent of tetraploid interphase nuclei in response to PDS in WT, *Brca1*$^{\Delta 11/\Delta 11}$, and CH12 cells. Error bars show the SEM from three independent experiments. Statistics: ****p < 0.001, ***p < 0.005  and **p < 0.01 comparing PDS doses.

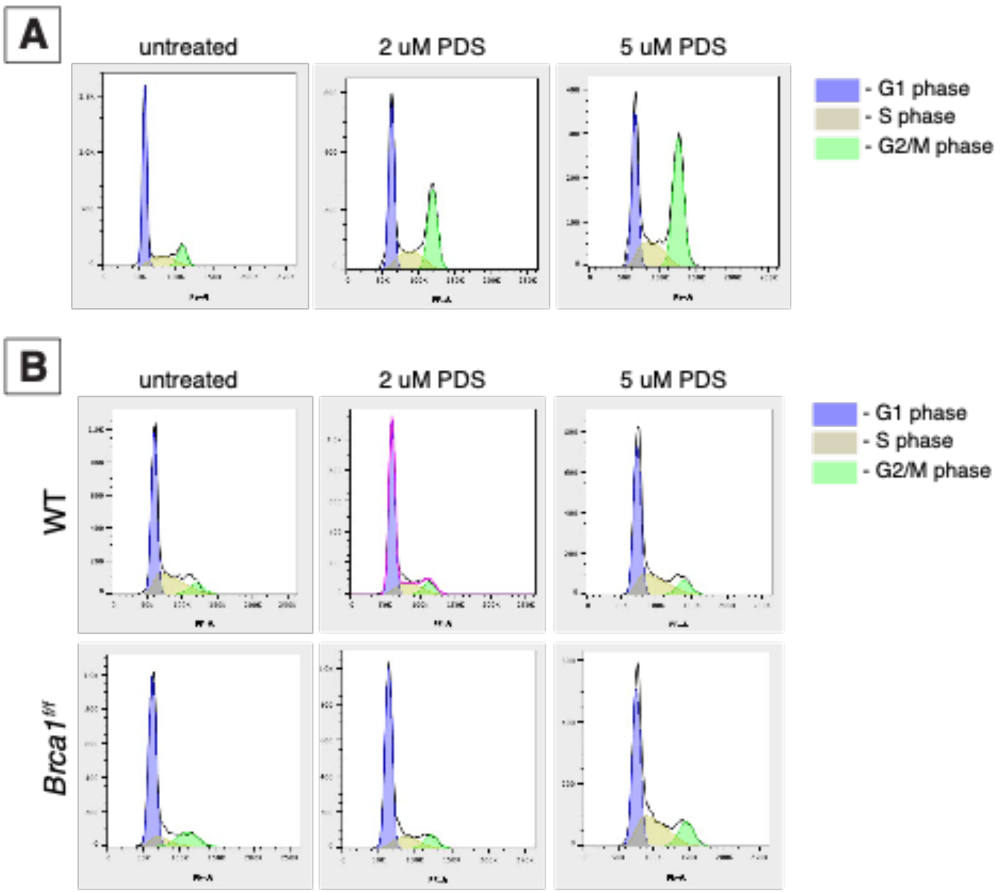

Fig. S4. **PDS-induced tetraploidy in primary B cells is associated with absence of G2/M arrest. A)** Representative flow cytometry histograms of PDS-treated CH12 cells assigned to different cell cycle stages (G1 – highlighted in blue, S – in yellow, G2/M – in green), based on the intensity of PI-staining (reflecting DNA content). **B)** Representative flow cytometry histograms of PDS-treated WT and *Brca1*$^{\Delta 11/\Delta 11}$ primary B cells assigned to different cell cycle stages (G1 – highlighted in blue, S – in yellow, G2/M – in green), based on the intensity of PI-staining (reflecting DNA content).

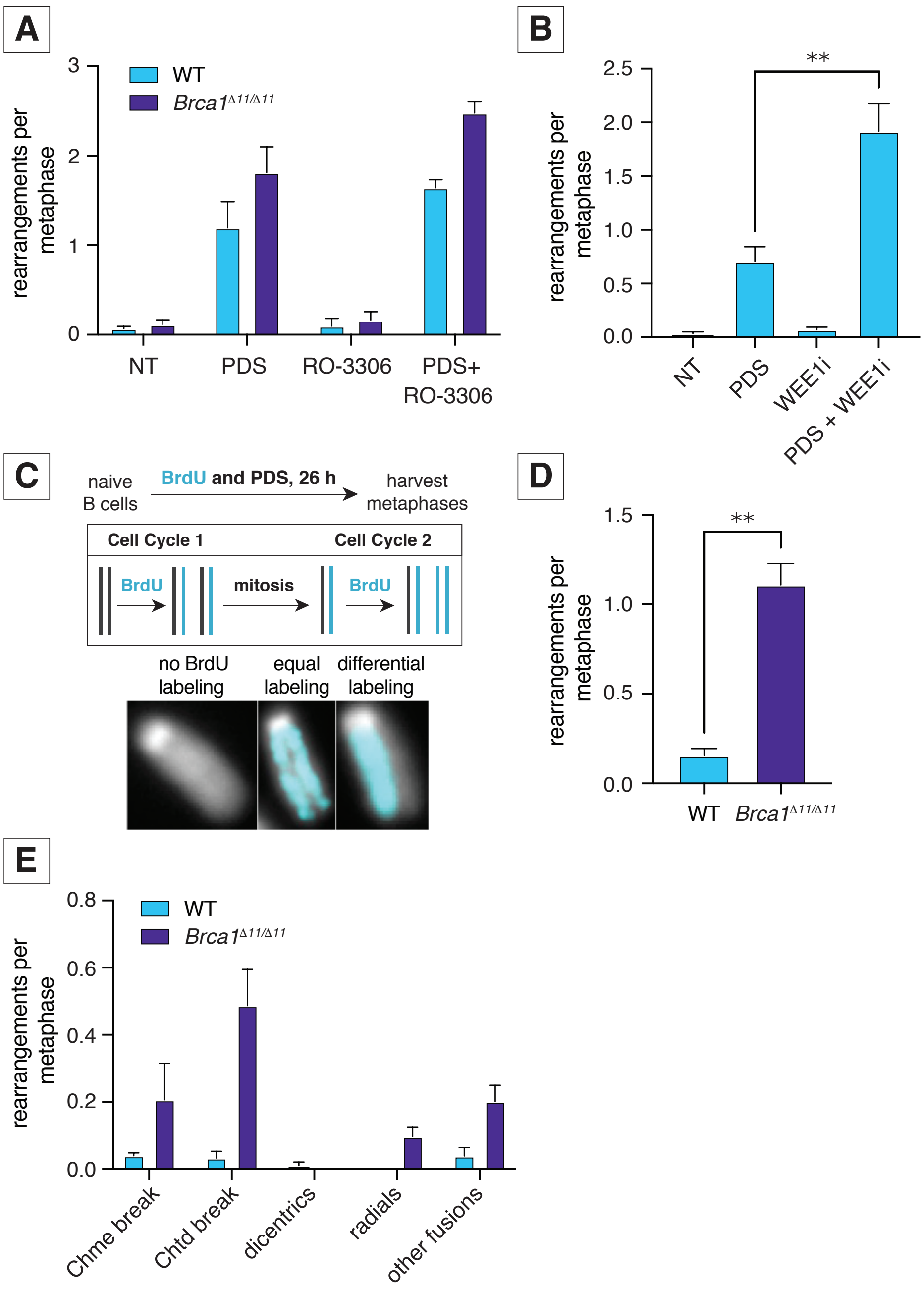

Fig. S5. **PDS treatment during a single cell cycle does not cause tetraploidy or extensive dicentric chromosome formation. A)** Number of DNA aberrations per metaphase in response to PDS combined with RO-3306 in WT and *Brca1*$^{\Delta11/\Delta11}$ cells. Error bars show the SEM from three independent experiments. **B)** Number of DNA aberrations per metaphase in response to PDS combined with WEE1i (AZD1775) in CH12 cells. Error bars show the SEM from three independent experiments. Statistics: $^{**}p < 0.01$ comparing 5 µM PDS and combination of 0.05 µM WEE1i with 5 µM PDS. **C)** Scheme of the experiment: Isolated from spleen naïve B cells were plated and stimulated accordingly to the protocol. Precisely at the plating time, 5 µM PDS and BrdU was added. BrdU labeling status was used to define metaphase spreads used in analysis: equally labeled BrdU cells were described as undergoing only one cell cycle in presence of PDS, while differentially labeled cells were defined as undergoing two (or more) cell cycles in the presence of PDS. **D)** Number of DNA aberrations per metaphase in response to 5 µM PDS during only one cell cycle in WT and *Brca1*$^{\Delta11/\Delta11}$ cells. Error bars show the SEM from three independent experiments. Statistics: $^{**}p < 0.01$ comparing WT and *Brca1*$^{\Delta 11/\Delta 11}$ cells exposed to 5 μM PDS. **E)** Different PDS-induced chromosomal rearrangements per plate during only one cell cycle in WT and *Brca1*$^{\Delta 11/\Delta 11}$ B cells exposed to 5 μM PDS. Error bars show the SEM from three independent experiments.

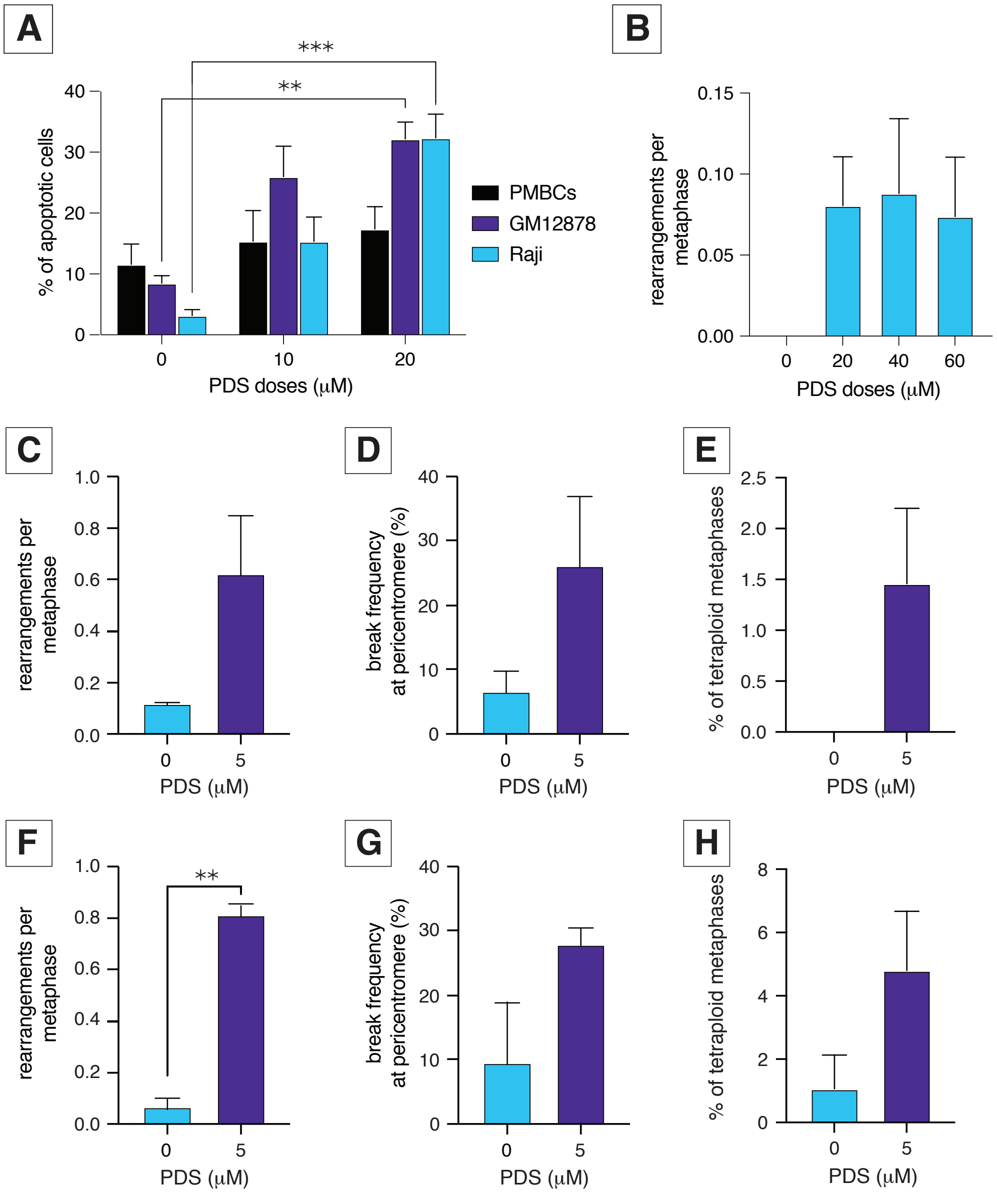

Fig. S6. **PDS causes pericentromeric breakage in human cancer cell lines. A)** Percent of TUNEL-positive apoptotic PBMCs, GM12878 and Raji cells in response to PDS treatment. Statistics: ***p < 0.005 and **p < 0.01 comparing PDS doses. **B)** Number of DNA aberrations per metaphase in response to high doses of PDS in PMBCs. Error bars show SEM from three independent experiments. **C)** Number of DNA aberrations per metaphase in response to PDS in U2OS cells. Error bars show SEM from three independent experiments. **D)** Frequency of pericentromeric breaks in response to PDS in U2OS cells. Error bars show SEM from three independent experiments. **E)** Percent of tetraploid metaphases in response to PDS in U2OS cells. Error bars show SEM from three independent experiments. **F)** Number of DNA aberrations per metaphase in response to PDS in MCF7 cells. Error bars show SEM from three independent experiments. Statistics: **p < 0.01 comparing PDS doses. **G)** Frequency of pericentromeric breaks in response to PDS in MCF7 cells. Error bars show SEM from three independent experiments. **H)** Percent of tetraploid metaphases in response to PDS in MCF7 cells. Error bars show SEM from three independent experiments.

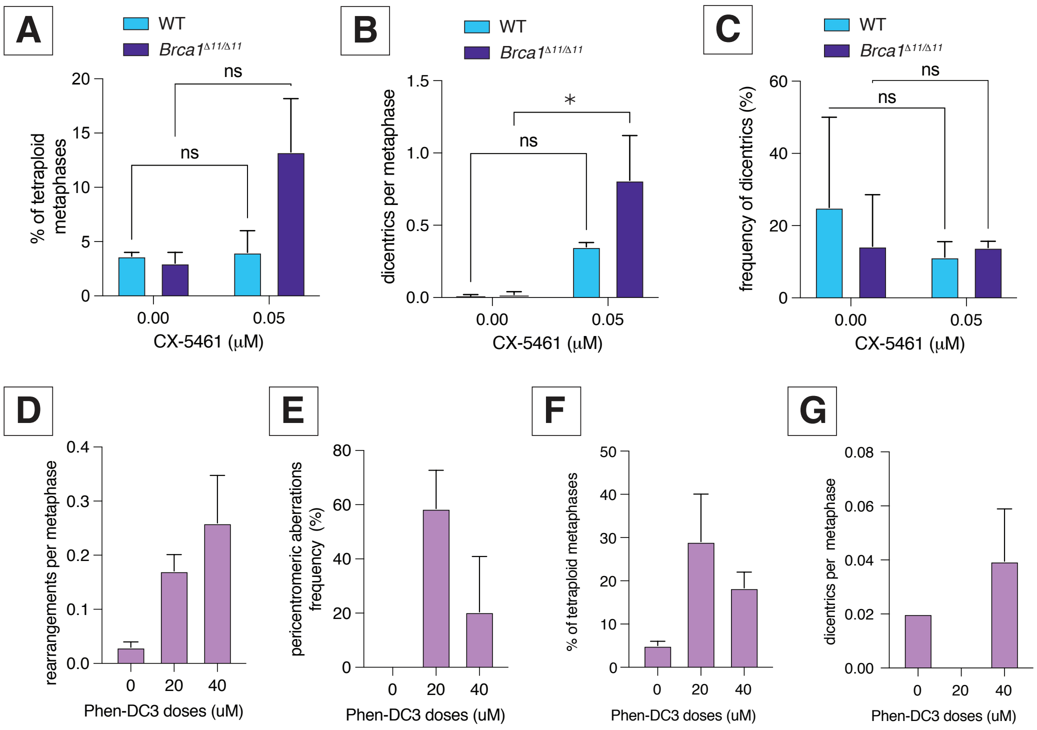

Fig. S7. **CX-5461- and Phen-DC3-induced DNA damage in primary B cells. A)** Percent of tetraploid metaphases in response to CX-5461 in WT and *Brca1*$^{\Delta 11/\Delta 11}$. Error bars show SEM from three independent experiments. **B)** Number of dicentric chromosomes per metaphase in response to CX-5461 in WT and *Brca1*$^{\Delta 11/\Delta 11}$ cells. Error bars show the SEM from three independent experiments. Statistics: *$p < 0.05$ comparing CX-5461 doses. **C)** Dicentric chromosome frequency in response to CX-5461 in WT and *Brca1*$^{\Delta 11/\Delta 11}$ cells. Error bars show the SEM from three independent experiments. **D)** Number of DNA aberrations per metaphase in response to Phen-DC3 in WT cells. Error bars show SEM from two independent experiments. **E)** Frequency of pericentromeric breaks in response to Phen-DC3 in WT cells. Error bars show SEM from two independent experiments. **F)** Percent of tetraploid metaphases in response to Phen-DC3 in WT cells. Error bars show SEM from two independent experiments. **G)** Number of dicentric chromosomes per metaphase in response to Phen-DC3 in WT cells. Error bars show the SEM from two independent experiments.

### Table S1. G4 analyses of centromere and rDNA sequences.

| | | | G4 hunter data | | | | | QGRS mapper | | |
|---|---|---|---|---|---|---|---|---|---|---|
| | | | | | | adjusted | | # of G4s (non- | | |
| sequence | length | %GC | G4 frequenc | # of G4s | score (highest | parameters* | score | overlapping) | score (highest) | **GenBank Accession** |
| mouse minor sa | 363 | 33.6 | 0 | 0 | na | 0 | | 0 | na | Z22170 |
| reverse complement | | | | | | | | 0 | na | |
| mouse major sa | 471 | 36.3 | 0 | 0 | na | 1 | 0.53 | 2 | 10 | M17407 |
| reverse complement | | | | | | | | 0 | na | |
| mouse 45s total | 45306 | 48.9 | 4.3 | 196 | 1.623 | from -1.484 to 1.623 | | 255 | 63 | BK000964 |
| | | | | | | | | 286 | 84 | |
| human rDNA tot | 42999 | 58.4 | 7 | 302 | 1.759 | from -1.692 to 1.759 | | 334 | 63 | U13369.1 |
| human chr 13 centromere repeats | | | | | | | | | | **T2T CHM13v2.0/hs1 Location** |
| other sat 13_27 | 138644 | 60.6 | 7.8 | 1086 | 1.519 | | | 1166 | 40 | chr13:12301367-12440010 |
| sat3_13_15 | 71663 | 46.1 | 2.4 | 169 | -1.385 | | | 2 | 15 | chr13:13172021-13243683 |
| bsat_13_8 | 7677 | 50.9 | 5.3 | 41 | -1.635 | | | 55 | 40 | chr13:11701524-11709200 |
| human chr 1 centromere repeats | | | | | | | | | | |
| beta | 496203 | 52.4 | 2.3 | 1145 | 1.837 | | | 2490 | 63 | chr1:128098616-128594818 |
| ct 1_7* | 39386 | 41.5 | 1.1 | 45 | -1.397 | | | 144 | 42 | chr1:127715078-127754463 |
| hsat2 | 592436 | 36.3 | 0 | 0 | na | | | 1 | 3 | chr1:126907666-127500101 |
| hsat3 | 214976 | 40.4 | 0.2 | 40 | 1.222 | | | 4595 | 21 | chr1:127500102-127715077 |
| hsat5* | 2130 | 59.2 | 7.5 | 16 | -1.269 | | | 33 | 30 | chr1:127881256-127883385 |
| censat1_17 | 2904 | 66.8 | 29.3 | 85 | 1.839 | | | 78 | 32 | chr1:128947980-128950883 |
| *transcribed | | | | | | | | | | |
| human LINE-1 | 5976 | 42.5 | 1 | 6 | 1.607 | | | 17 | 58 | M19503.1 |

G4 Hunter parameters Window size: 25, Threshold: 1.2 (default)
G4 Hunter adjusted paramete Window size: 40, Threshold: 0.5

QGRS parameters Search Parameters: QGRS Max Length: 30 | Min G-Group Size: 2 | Loop size: from 0 to 36

**Table S2. The frequency of dicentric chromosomes correlates with tetraploidy.** a) Mantel-Haenzel test for correlation between tetraploidy and elevated rate of dicentric chromosomes in response to 5 μM PDS in WT and *Brca1*$^{\Delta 11/\Delta 11}$ B cells. 3 independent experiments were performed.Null hypothesis: the elevated rate of dicentric chromosomes doesn't correlate with tetraploidy. Alternative hypothesis: the elevated rate of dicentric chromosomes correlates with tetraploidy. b) Mantel-Haenzel test for correlation between tetraploidy and appearance more than one dicentric chromosome in the cell in response to 5 μM PDS in WT and *Brca1*$^{\Delta 11/\Delta 11}$ B cells. 3 independent experiments were performed. Null hypothesis: More than one dicentric chromosomes per cell doesn't correlate with tetraploidy. Alternative hypothesis: More than one dicentric chromosomes per cell correlates with tetraploidy.

**a) The frequency of dicentrics does not correlate with tetraploidy**

**WT 5 uM PDS**

| | | |
|---|---|---|
| **continuity correction? enter 'y' or 'n':** | y | 0.5 |
| **chi-square:** | 27.037 | |
| **d.f.:** | 1 | |
| **P-value:** | 2.00E-7 | Null hypothesis is rejected |

| experiment | | diploids | tetraploids | | |
|---|---|---|---|---|---|
| 1 | no dicentrics | 208 | 17 | 3.10638298 | 0.78246335 |
| | dicentrics | 6 | 4 | | |
| 2 | no dicentrics | 183 | 17 | 0.56097561 | 0.39262344 |
| | dicentrics | 4 | 1 | | |
| 3 | no dicentrics | 200 | 24 | 2.41048035 | 0.51088993 |
| | dicentrics | 2 | 3 | | |

**Brca1 5 uM PDS**

| | | |
|---|---|---|
| **continuity correction? enter 'y' or 'n':** | y | 0.5 |
| **chi-square:** | 114.909 | |
| **d.f.:** | 1 | |
| **P-value:** | 8.24E-27 | Null hypothesis is rejected |

| experiment | | diploids | tetraploids | | |
|---|---|---|---|---|---|
| 1 | no dicentrics | 185 | 18 | 13.6919831 | 3.85299087 |
| | dicentrics | 15 | 19 | | |
| 2 | no dicentrics | 196 | 16 | 7.36326531 | 2.81156458 |
| | dicentrics | 22 | 11 | | |
| 3 | no dicentrics | 186 | 30 | 8.79183673 | 3.78249609 |
| | dicentrics | 15 | 14 | | |

**b) More than one dicentrics per plate does not correlate with tetraploidy**

**WT 5 uM PDS**

| | | |
|---|---|---|
| **continuity correction? enter 'y' or 'n':** | y | 0.5 |
| **chi-square:** | 4.11 | |
| **d.f.:** | 1 | |
| **P-value:** | 0.043 | Null hypothesis is rejected |

| experiment | | diploids | tetraploids | | |
|---|---|---|---|---|---|
| 1 | one dicentric | 5 | 2 | 0.8 | 0.56 |
| | more than one dicentric | 1 | 2 | | |
| 2 | one dicentric | 4 | 1 | 0.66666667 | 0.22222222 |
| | more than one dicentric | 0 | 1 | | |
| 3 | one dicentric | 2 | 0 | 1.2 | 0.36 |
| | more than one dicentric | 0 | 3 | | |

**Brca1 5 uM PDS**

| | | |
|---|---|---|
| **continuity correction? enter 'y' or 'n':** | y | 0.5 |
| **chi-square:** | 27.48 | |
| **d.f.:** | 1 | |
| **P-value:** | 1.59E-7 | Null hypothesis is rejected |

| experiment | | diploids | tetraploids | | |
|---|---|---|---|---|---|
| 1 | one dicentric | 15 | 8 | 4.85294118 | 1.89013841 |
| | more than one dicentric | 0 | 11 | | |
| 2 | one dicentric | 20 | 5 | 3.33333333 | 1.38888889 |
| | more than one dicentric | 2 | 6 | | |
| 3 | one dicentric | 15 | 7 | 3.62068966 | 1.37336504 |
| | more than one dicentric | 0 | 7 | | |
